# Tetraspanin CD82 maintains HTLV-1 biofilm polarization and is required for efficient viral transmission

**DOI:** 10.1101/2023.05.02.538526

**Authors:** Coline Arone, Samuel Martial, Julien Burlaud-Gaillard, Maria-Isabel Thoulouze, Philippe Roingeard, Hélène Dutartre, Delphine Muriaux

## Abstract

The human T-lymphotropic virus type-1 (HTLV-1) is an oncogenic retrovirus whose transmission relies primarily on cell-to-cell contacts as cell-free viruses are poorly infectious. Among the intercellular transmission routes described, HTLV-1 biofilms are adhesive structures polarized at the cell surface that confine virions in a protective environment, which is believed to promote their simultaneous delivery during infection. Here, we show that several tetraspanins are enriched in HTLV-1 biofilms and incorporated into the viral envelope. However, we report that only tetraspanin CD82 interacts with HTLV-1 Gag which initiates its polarization into viral biofilms. Also, we demonstrate that CD82 maintains HTLV-1 biofilm polarization and favors viral transmission, as its silencing induces a complete reorganization of viral clusters at the cell surface and reduces the ability of infected T-cells to transmit the virus. Our results highlight the crucial role of CD82 in the architectural organization of HTLV-1 biofilms and their transfer through intercellular contacts.

## INTRODUCTION

The Human T-Lymphotropic virus type-I (HTLV-1) is the most potent oncogenic virus known to date, with a minimal estimation of 5 to 10 million people infected worldwide^1,2^. Although most people remain asymptomatic following HTLV-1 exposure, chronic infections can lead to aggressive pathologies with poor prognoses such as adult T-cell Leukemia (ATL)^3,4^, progressive inflammatory disorders like HTLV-1-associated myelopathy/tropical spastic paraparesis (HAM/TSP)^5,6^, and other less severe pathologies (uveitis, dermatitis, myositis, etc.)^7^. *In vivo*, HTLV-1 is primarily detected in CD4(+) T-cells and to a lesser extent in CD8(+) T-cells, monocytes, or dendritic cells. Transmission among individuals occurs through three main routes: mother to child during breastfeeding, sexual contact, and exposure to HTLV-1-infected blood products^8^. Overall, this virus represents a major health issue as no therapeutic strategy allows the efficient protection of exposed individuals. Therefore, fundamental research on HTLV-1 is essential to promote the expansion of diagnosis methods and pertinent therapeutic approaches. HTLV-1 belongs to the Deltaretrovirus genus of the Retroviridae family and thus requires the expression of its main structural protein, the Gag polyprotein, to drive particle assembly, budding, and release from the plasma membrane^9^. In terms of structure, HTLV-1 is enveloped and contains two single-stranded genomic RNA molecules sheltered by an icosahedral capsid^10^. As such, viral particles are mostly spherical and display sizes ranging from 76 to 175 nm in diameter^11,12^. Although HTLV-1 structural details have been uncovered along with its macroscopic transmission routes, HTLV-1 cell-to-cell transmission pathways remain to be completely elucidated. Unlike most retroviruses, evidence strongly suggests that cell-free virions are poorly infectious *in vivo*. This belief originated from transfusion studies showing that, unlike plasma fractions or plasma derivatives where viral RNAs were rarely detected, cellular blood components were strictly needed for productive infections^13,14^. *In vitro*, previous experiments supported this notion and demonstrated that cell-free viruses do not efficiently infect primary T-lymphocytes as compared to co-cultures with HTLV-1-infected cell lines^15,16^.

HTLV-1 cell-to-cell transmission strategies described in models mimicking its preferential tropism (*i.e.*, CD4(+) T-cells) include the virological synapse, the cellular conduits, and viral biofilms (for a review see^17^). These three pathways, which may not be mutually exclusive, display similarities as they all allow the transfer of viral particles to target cells and involve plasma membrane remodeling. On one hand, the virological synapse is described as a cytoskeleton-dependent, stable adhesive junction that enables direct virus transfer^18^. On the other hand, cellular conduits are membrane protrusions that act as an avenue for the movement of viral particles toward target cells^19^. At last, viral biofilms are cell-surface aggregates of viruses enriched in carbohydrates, glycoproteins (*i.e.*, agrin), intercellular adhesion molecules (*i.e.*, ICAM-1), and ECM components up-regulated by HTLV-1 (*i.e.*, galectin-3, collagen, fibronectin, sialyl Lewis X), that promote virions accumulation at one or several poles of the infected cells^20–24^. These structures allow the bulk delivery of infectious particles to target cells and may help to protect HTLV-1 virions from immune attacks^25,26^. In addition to chronically infected T-cells, HTLV-1 biofilms were also observed at the surface of primary CD4(+) T-cells from HAM/TSP patients or asymptomatic HTLV-1 carriers^24^. While the relative importance of the different cell-to-cell transmission routes is difficult to ascertain *in vivo*, the removal of biofilms by heparin washes reduces the infectious capacity of HTLV-1-producing cells by 80% *in vitro*^24^. In addition, biofilms detached from HTLV-1 infected cells are infectious *in vitro,* as opposed to cell-free virions released in the supernatant of the same cells^16,24^, indicating that the transfer of HTLV-1 biofilms through cell-to-cell contacts is the most efficient pathway to infect new cells.

To identify molecular actors involved in HTLV-1 biofilm architecture and transmission, we focused here on the role of specific biofilm components belonging to the Transmembrane 4 Superfamily (TM4SF) of proteins, also known as tetraspanins. Tetraspanins are four-span transmembrane proteins organized in tetraspanin-enriched microdomains (TEMs) at the plasma membrane, where they self-associate through their large extracellular loops or interact with various protein complexes, thereby generating a hierarchical network of interactions (for a review see^27^). They are expressed by all metazoans, with 33 members in mammals, including CD9, CD63, CD81, CD82, and CD151. Among their known functions, tetraspanins regulate T-cells adhesion (cell-to-cell and cell-to-ECM), synapse formation, cell motility, and membrane compartmentalization. In addition, several pathogens hijack these surface proteins for infection, with CD81 being the main receptor of hepatitis C virus entry for instance, or CD151 being involved in human papillomavirus endocytosis^28,29^. Concerning retroviruses, it has been shown for the human immunodeficiency virus 1 (HIV-1) that viral particles accumulate in membrane microdomains enriched in CD63, CD81, and CD82 in infected cells^30–32^. In addition, CD81, which is found in the viral membrane, is required for the polarization of HIV-1 Gag clusters at the T-cell surface^30^. For HTLV-1, Mazurov and collaborators demonstrated that CD9, CD53, CD63, and CD82 are found in viral aggregates at the surface of chronically infected T-cells, although no function was addressed in their study. Nevertheless, it was shown that HTLV-1 Gag can form intracellular complexes with CD81 and CD82 intracellular loops when over-expressed in HEK293 cells^21,33^. Another study performed with chronically infected-T cells reported the interaction between CD82 and HTLV-1 envelope glycoprotein (Env) which remain associated during their trafficking to the plasma membrane through the secretory pathway^34^. Overall, these observations suggest a function for cellular tetraspanins in viral proteins’ trafficking, viral assembly, and/or biofilm formation, which could subsequently contribute to HTLV-1 cell-to-cell transmission.

Here, we asked whether these tetraspanins could be incorporated into viral particles within HTLV-1 biofilms, participate in biofilm biogenesis at the T-cell surface, and play a role in the subsequent transmission of HTLV-1 biofilms to target T-cells. In our study, we show the enrichment of CD9, CD81, and CD82 in HTLV-1 biofilms released from infected T-cells using mass spectrometry coupled to immunoblotting, along with the strong polarization of these three transmembrane molecules into cell-attached biofilms using immunofluorescence microscopy. Accordingly, we demonstrate the incorporation of CD9, CD81, and CD82 in the viral lipidic envelope using super-resolution STED microscopy and immuno-electron microscopy. At last, using shRNAs to silence the expression of CD9, CD81, or CD82 in chronically infected T-cells, we identify a crucial role for CD82 in the regulation of HTLV-1 biofilms polarization at the cell surface. Its silencing depolarizes HTLV-1 Env(+) aggregates and reduces the ability of HTLV-1-producing cells to infect target T-cells. Overall, our results highlight a specific role for the tetraspanin CD82 in HTLV-1 clustering and biofilm polarization, which we document as two essential parameters for efficient cell-to-cell transmission.

## RESULTS

### Super-resolution microscopy imaging confirms the integrity of HTLV-1 biofilms isolated from infected T-cells

Viral biofilms isolated from HTLV-1-infected T-cells are known to be infectious units as they can productively infect primary T-cells and monocyte-derived dendritic cells *in vitro*^16^. Thus, they are relevant minimal systems to study molecular components linked to HTLV-1 cell-to-cell transmission. In the present study, we produced and purified non-fluorescent and fluorescent HTLV-1 biofilms to further investigate their structural (Figure 1) and molecular composition (Figure 2).

**Figure 1:**
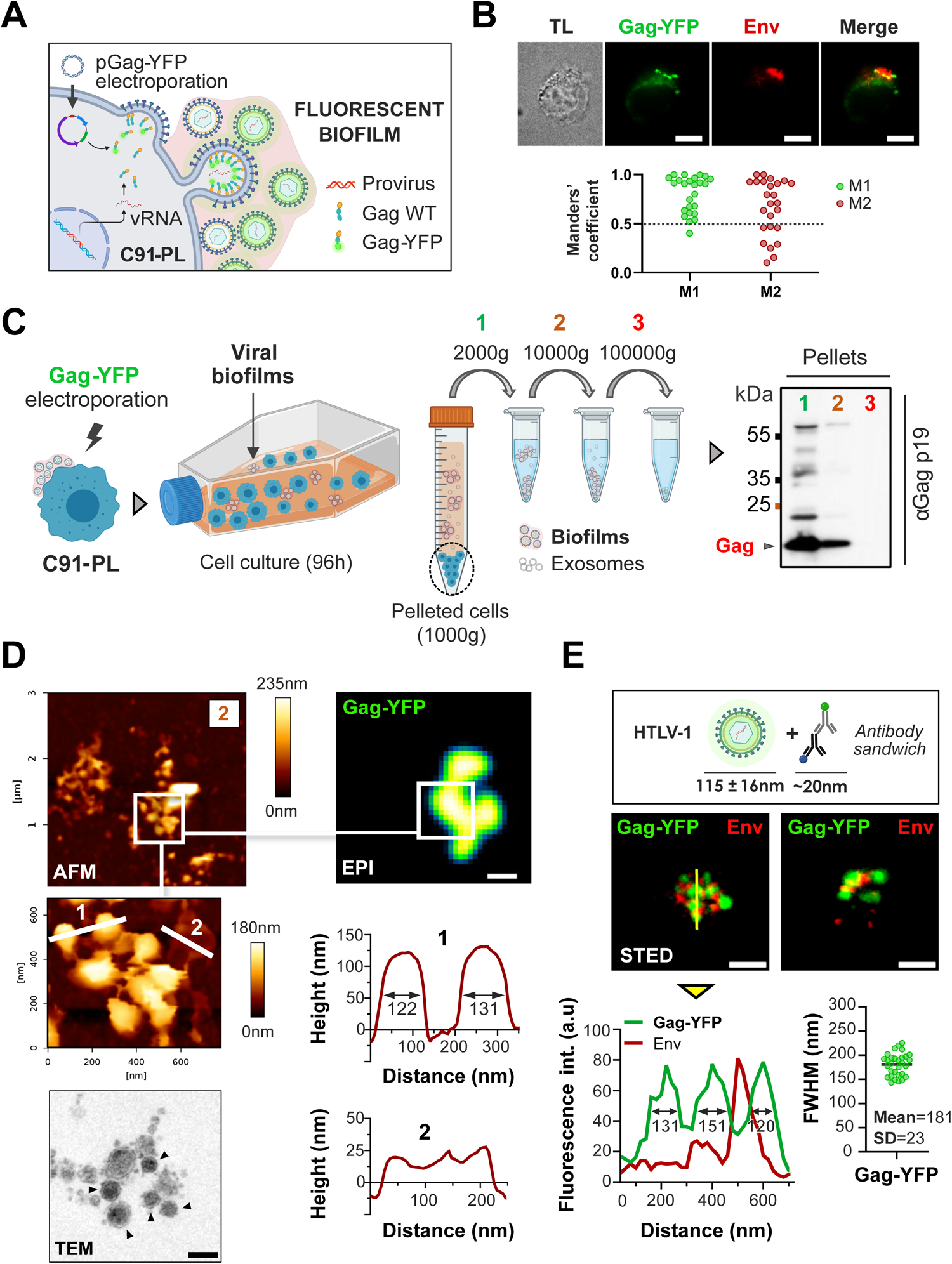
Super-resolution microscopy imaging confirms the integrity of HTLV-1 biofilms isolated from infected T-cells. **(A)** Scheme: Electroporation of HTLV-1 chronically infected T-cells (C91-PL) with a plasmid encoding HTLV-1 Gag-YFP. **(B)** Top: Immunofluorescence images show the successful incorporation of Gag-YFP (green) into neo-formed Env(+) virions (red). TL: Transmitted light. Below: Manders’ coefficients as a measure of colocalization between (n = 25 cells): 0 = non-overlapping images; 1 = 100% co-localization between both channels. M1 is the fraction of Gag overlapping Env and M2 is the fraction of Env overlapping Gag. Scale bars = 10 µm **(C)** Left: Protocol for isolation of fluorescent biofilms from chronically infected T-cells (C91-PL). Right: Immunoblot probed for HTLV-1 Gagp19 showing Gag expression in biofilms isolated at 2000g (1), 10000g (2), or 100000g (3). **(D)** Top: Atomic force microscopy correlated to fluorescence imaging of native Gag-YFP(+) biofilms pelleted at 10000g (2). Gag-YFP(+) structures (biofilms) revealed by wide-field fluorescence microscopy were imaged by AFM (topographic images). Higher magnification shows spherical particles whose size corresponds to HTLV-1 diameter (FWHM on the 1^st^ cross-section/upper plot = 122nm and 131nm) and thin structures bound to the particles (2^nd^ cross-section/lower plot). Scale bar = 500nm. Bottom left: Transmission electron microscopy image of cell-free HTLV-1 biofilm. Individual viral particles are indicated with black arrows. Scale bar = 200nm. **(E)** Top: Scheme of HTLV-1 Gag-YFP(+) particles’ expected size after staining. Middle: Representative STED microscopy images of fixed Gag-YFP(+) biofilms pelleted at 10000g and stained for the viral envelope protein (Env). Scale bars = 500nm. Bottom: The yellow cross-section is plotted below the corresponding image (left graph) where the three green peaks correspond to individual particles (FWHM = 131nm, 151nm, and 120nm). The right plot shows the FWHM distribution of n = 25 Gag-YFP(+) particles. FWHM: full width at half maximum. SD: standard deviation of the mean.

To first decipher the structure and morphology of HTLV-1 biofilms, C91-PL chronically infected T-cells, which produce large and polarized viral biofilms, were electroporated with a plasmid encoding the fluorescent viral protein Gag-YFP (Figure 1A). Of note, the fusion of the YFP tag at the C-terminal end of Gag was shown to not impair its trafficking or pseudo-particles’ assembly^35^. Twenty hours post-electroporation, living Gag-YFP(+) infected cells started to exhibit viral aggregates that accumulated within a few hours into large, polarized biofilms following lateral movements of preformed Gag-YFP(+) clusters toward intercellular contacts (supplemental video 1 and Figure S1). These Gag-YFP(+) biofilms colocalized with viral envelope proteins (Figure 1B), which confirms that neo-formed Env(+) particles successfully incorporate Gag-YFP molecules. Then, we maintained Gag-YFP(+) C91-PL cells at a high cellular density to promote the release of biofilms in the culture supernatant that were collected using sequential centrifugations (2000g, 10000g, and 100000g), as shown in Figure 1C. While cell-free enveloped viruses are usually sedimented by >100000g ultracentrifugation^36^, most HTLV-1 particles detected using Gag-p19 immunoblotting were sedimented at 10000g, suggesting their nearly complete retention in dense structures corresponding to viral biofilms (Figure 1C). Based on these observations, we always harvested HTLV-1 biofilms at 10000g throughout the study.

To assess whether intact HTLV-1 biofilms were successfully isolated using this method, we imaged our samples using an atomic force microscope (AFM) coupled with fluorescence microscopy, which allowed us to go beyond the light diffraction limit and distinguish individual virions (Figure 1D). As shown in the magnified AFM image in Figure 1D, Gag-YFP(+) biofilms contained spherical particles with a mean central height of 126nm□±□8nm (*e.g*., 1st cross-section). This particle size is consistent with the diameter of HTLV-1 virions measured by cryogenic transmission electron microscopy^11,12^ (*i.e.*,115nm ± 16nm, with a broad size distribution ranging from 76 to 175 nm). Moreover, Gag-YFP(+) viral particles appeared interconnected by a network of fine filamentous structures (Figure 1D) that exhibited a mean height of 29nm□±□6nm (*e.g.*, 2nd cross-section). Although these structures are unlikely to be plasma membrane extensions since they are five times thicker than model lipid bilayers^37^, they could nonetheless correspond to scaffolds of extracellular matrix (ECM) components. This hypothesis is supported by transmission electron microscopy images of cell-free HTLV-1 biofilms where viral particles are associated with electron-dense ECM structures (Figure 1D).

Following the same isolation protocol, we explored if these fluorescent particles successfully incorporated the Env glycoprotein using super-resolution stimulated emission-depletion (STED) microscopy. Most Gag-YFP(+) particles (full width at half maximum = 181nm□±□23nm) were positive for Env and clustered together in biofilms (Figure 1E). Although some particles did not exhibit Env signals at their surface, it is known that HTLV-1 particles contain variable numbers of Env molecules, most of which are unevenly distributed in the viral lipidic envelope^38^. Overall, our results demonstrate the successful isolation of HTLV-1 biofilms containing unaltered Gag(+)/Env(+) viral particles.

### HTLV-1 biofilms released from infected T-cells are enriched in tetraspanins

To screen for key molecular components of HTLV-1 biofilms, we performed a large-scale identification of viral and cellular proteins contained in these isolated biofilms using mass spectrometry. Biofilms produced from C91-PL chronically infected cells were isolated (as in Figure 1C), inactivated, and processed on a mass spectrometer (Figure 2A). Among all the proteins identified, we excluded contaminants (including mitochondrial and ribosomal proteins) and considered only the 100 most abundant proteins (based on their IBAQ score) that were identified with at least three peptides in three independent replicates. As expected, viral proteins (Gag, Env) were detected in C91-PL biofilms (Figure 2B), along with proteins up-regulated by HTLV-1 infection (*e.g.*, Fascin^39^, Vimentin, GSE17718 database^40^) and extracellular matrix proteins like galectins^41^, which are known components of HTLV-1 biofilms^24^ (Figure 2B). Thus, the presence of HTLV-1-associated proteins validated the successful isolation of viral biofilms, allowing the reliable identification of new biofilm components.

**Figure 2:**
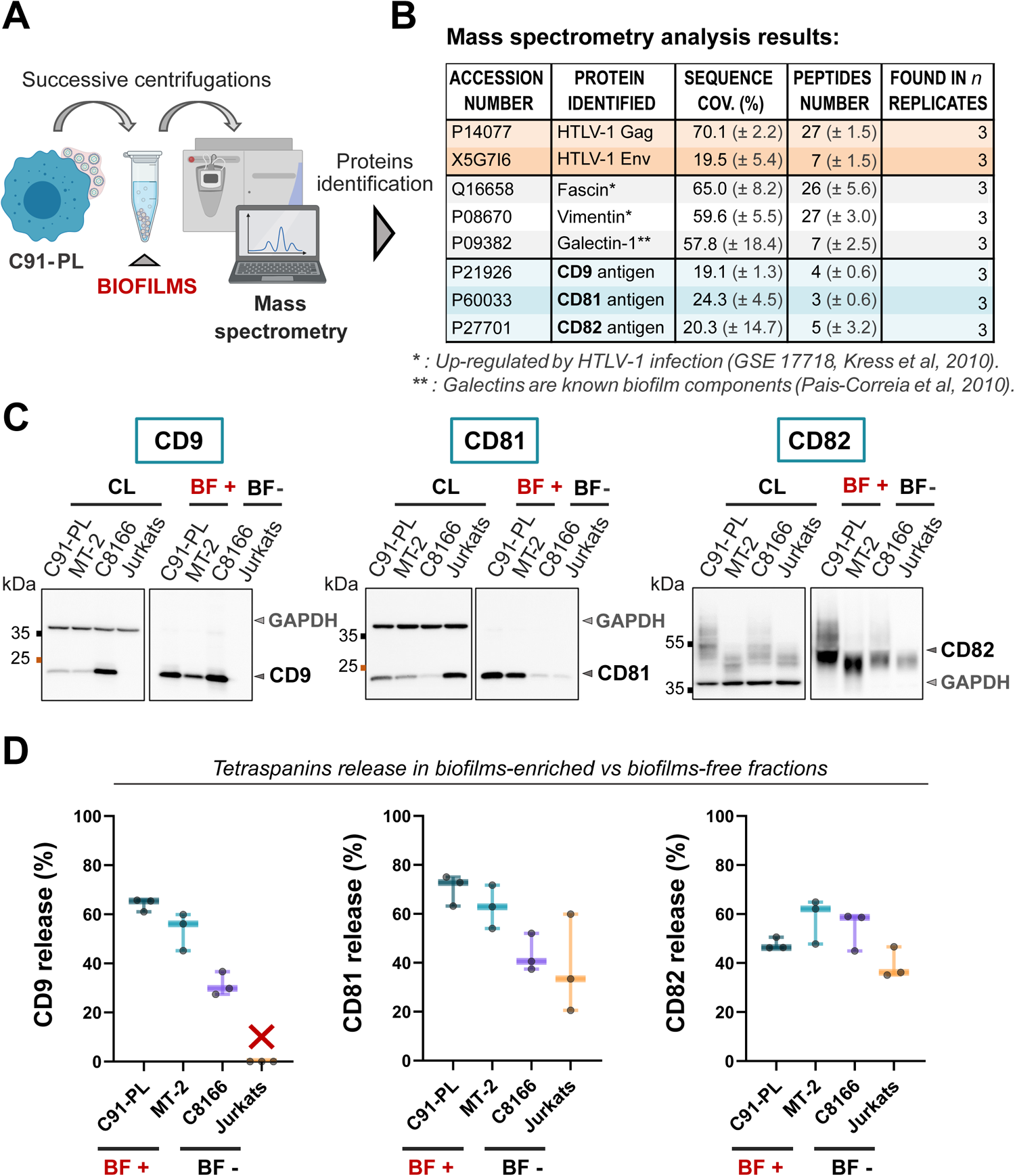
CD9, CD81, and CD82 are detected and enriched in viral biofilms released from HTLV-1 chronically infected T-cells. **(A)** Summary of the protocol used to isolate wild-type biofilms released in the cell-culture medium of HTLV-1 chronically infected T-cells (C91-PL). This protocol is detailed in Figure 1C. **(B)** Table of proteins identified with high confidence by mass spectrometry (MS) in the biofilms-enriched supernatant of HTLV-1 infected T-cells (C91-PL). The complete list of proteins identified with the highest confidence by MS is provided in Supplemental Figure 2. Along with viral proteins (orange) and cellular proteins (grey) known to be up-regulated by HTLV-1 infection (*) or present in biofilms (**); CD9, CD81, and CD82 tetraspanins (blue) were identified in biofilms-enriched fractions. Results were obtained from three independent experiments; standard deviation values are given in brackets. **(C)** Immunoblots of CD9, CD81, and CD82 contained in cell lysates (CL); biofilms-enriched supernatants (BF +); or biofilms-free supernatants (BF -) from four different cell lines: HTLV-1 infected cells producing biofilms (C91-PL, MT-2); HTLV-1 infected cells deficient in viral particles production (C8166); or non-infected T-cells (Jurkats). Endogenous GAPDH is used as a loading control for cell lysates. **(D)** Tetraspanins release in biofilms was calculated as the ratio of the amount of tetraspanins in BF (corrected by the dilution factor DF) to the total amount of tetraspanins (CL + BF) following this formula: ((BF*DF) / (BF*DF + CL*DF))*100. Box plots show the corresponding results for CD9, CD81, and CD82 released by each cell line in %. CD9 was not detected in Jurkat cells lysate or supernatant as shown by the red cross. Results were obtained from three independent experiments.

Among the 100 most abundant proteins detected in isolated biofilms, 69 were found to be involved in protein-protein interaction networks related to the cytoskeleton or the cellular surface (supplemental Figure S2). Importantly, a cluster of 3 tetraspanins was identified with high confidence in HTLV-1 biofilms: CD9, CD81, and CD82 (Figure 2B). To further determine whether CD9, CD81, and CD82 were enriched specifically in HTLV-1 biofilms, we quantified their release in the supernatant of four different cell lines (Figure 2C): HTLV-1 infected T-cells producing biofilms (C91-PL, MT-2); HTLV-1 infected T-cells expressing viral accessory proteins but deficient in virions production (C8166); or non-infected control T-cells (Jurkat). Tetraspanins release was calculated as the ratio of the amount of tetraspanins in biofilms-enriched fractions (isolated from C91-PL/MT-2 T-cells; *BF+*) or biofilms-free fractions (isolated from C8166/Jurkat T-cells; *BF-*) to the total amount of tetraspanins (see “Western blot analysis” in the materials and methods section). As shown in Figure 2D, around 60% of total CD9 was detected in BF(+) fractions from C91-PL/MT-2 cells against less than 35% in BF(-) fractions from C8166 cells or Jurkat cells, where CD9 expression was below detection levels. Although CD81 was detected in BF(-) fractions from control cells, it was similarly enriched in BF(+) fractions from HTLV-1-producing T-cells (Figure 2D). At last, CD82 release was equivalent in cells producing biofilms (C91-PL, MT-2) and infected cells deficient for viral particles’ production (C8166) (Figure 2D). Nevertheless, CD82 was enriched in the supernatant of infected T-cells (C91-PL, MT-2, C8166) as compared to non-infected T-cells (Jurkat), similar to CD9 and CD81, suggesting that the expression and release of these tetraspanins could be modulated by the infection. Altogether, these results highlight the preferential enrichment of several tetraspanins in HTLV-1 biofilms released from infected T-cells.

### CD9, CD81, and CD82 are polarized in biofilms at the surface of HTLV-1 producing T-cells

Following the detection of CD9, CD81, and CD82 in released HTLV-1 biofilms, we examined their localization at the surface of chronically infected T-cells (C91-PL), compared to infected T-cells that do not produce viral particles (C8166) or non-infected T-cells (Jurkat). Accordingly, fixed C91-PL, C8166, and Jurkat cells were immuno-stained for CD9, CD81, or CD82 and imaged on a confocal microscope (Figure 3A). Two major phenotypes emerged for the three molecules: a “polarized” pattern where tetraspanins were clustered on one side of the cell surface, mainly observed in HTLV-1-producing cells (C91-PL); and a “dispersed” pattern where their signal was distributed all over the cell periphery, mainly observed in non-producing infected cells (C8166) or non-infected control cells (Jurkat) (see supplemental Figure S3A for a more detailed image gallery). We quantified the occurrence of these phenotypes per cell line in Figure 3B. While 70% to 90% of control cells displayed a dispersed pattern for all tetraspanins tested, this phenotype was the opposite in C91-PL cells (Figure 3B), where at least 70% of cells carried polarized aggregates of CD9, CD81, and CD82. To check if viral assembly was the minimal requirement to cluster CD9, CD81, and CD82 at the cell surface, C8166 cells were electroporated with HTLV-1 Gag-YFP constructs (Figure 3C), knowing that Gag is the minimal system for pseudo-viral particles assembly and release^35^. This resulted in a small increase of cells displaying a polarized pattern for CD9 (from 13% to 33%), and a completely reversed phenotype for both CD81 and CD82 that were found polarized in more than 60% of Gag-YFP(+) C8166 cells (Figure 3D, see supplemental Figure S3B for a more detailed image gallery). This indicates that (i) HTLV-1 Gag is required but not sufficient to induce an extensive CD9 polarization in cells lacking other viral assembly components (*i.e*, Env), whereas (ii) Gag expression almost completely restores the polarized pattern observed for CD81 and CD82 in C91-PL cells.

**Figure 3:**
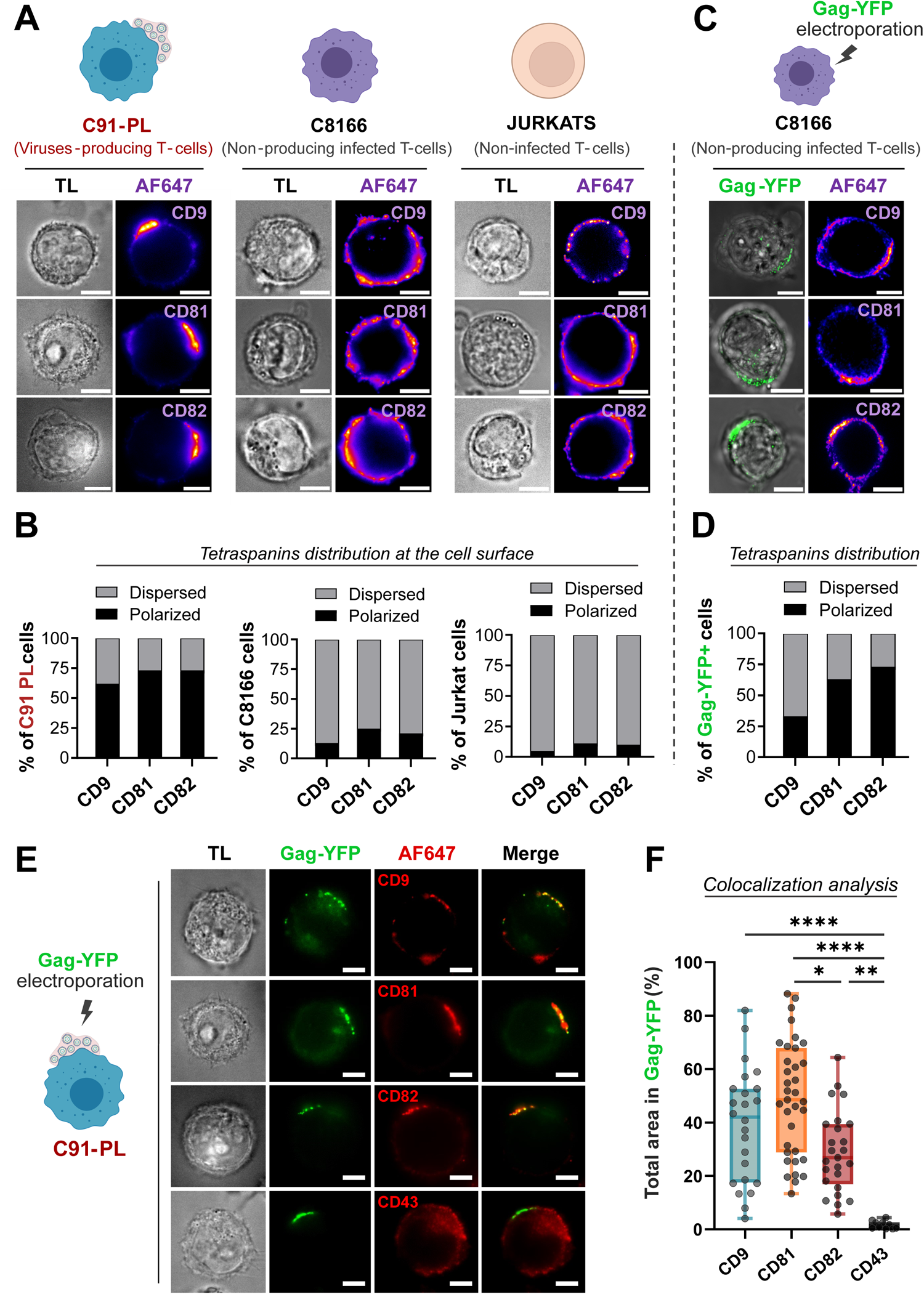
CD9, CD81, and CD82 are polarized in viral biofilms at the surface of HTLV-1 producing T-cells. **(A)** Representative immunofluorescence microscopy images showing CD9, CD81, and CD82 tetraspanins localization at the surface of fixed non-permeabilized cells. Two major patterns were observed in HTLV-1 chronically infected C91-PL cells (in red) and control cells (C8166 and Jurkat): “Polarized” – *Tetraspanins are clustered on one side of the cell surface*, or “Dispersed” – *Tetraspanins signal is distributed all over the cell periphery as punctuated small dots*. Representative images of the most abundant pattern (“polarized” or “dispersed”) observed in each condition are shown. TL: Transmitted light. Scale bars = 10µm. **(B)** Quantification of CD9, CD81, and CD82 distribution patterns in the cell population for each cell line (n > 30 cells counted for each condition). On the histograms, grey bars = % of cells displaying the dispersed pattern; black bars = % of cells displaying the polarized pattern for each tetraspanin tested. **(C)** Representative immunofluorescence microscopy images of tetraspanins localization at the surface of C8166 cells upon transient Gag-YFP expression. Scale bars = 10µm. **(D)** CD9, CD81, and CD82 distribution patterns in the cell population were quantified as in **(B)** (n > 30 cells counted for each condition) and reported on the histogram. **(E)** C91-PL cells were electroporated with Gag-YFP and stained for CD9, CD81, CD82, or CD43. Representative immunofluorescence microscopy of Gag-YFP(+) C91-PL cells showing the localization of CD9, CD81, CD82, or CD43 (red) along with Gag-YFP(+) biofilms at the cell surface (green). Scale bars = 10µm. **(F)** Colocalization analysis: Box plot showing the total area of CD9, CD81, CD82, or CD43 signals (Channel 2) overlapping Gag-YFP(+) biofilms (Channel 1) in %, quantified for each condition (n > 25 cells). See supplemental Figure S3C for thresholding and calculation details. Kruskal-Wallis statistical tests were used to assess any significant difference: * = p.value ≤ 0.05; ** = p.value ≤ 0.01; **** = p.value ≤ 0.0001.

To investigate whether these polarized tetraspanins were associated with HTLV-1 biofilms at the cell surface, we used chronically infected T-cells (C91-PL) electroporated with Gag-YFP constructs (schematized in Figure 1A). Gag-YFP(+) T-cells were fixed and stained for CD9, CD81, CD82, or control CD43. CD43 is a transmembrane glycoprotein reported to be distributed all around the cell periphery without being specifically enriched in HTLV-1 biofilms^42^ and is therefore used here as a negative control. As expected, confocal images showed no colocalization between Gag-YFP(+) biofilms and CD43 (Figure 3E). However, an extensive colocalization was observed between CD9, CD81, or CD82 and Gag-YFP(+) biofilms that were polarized together on one side of the cell surface (Figure 3E). To quantify the degree of colocalization, we calculated the percentage of the total area occupied by each tetraspanin that was overlapping with Gag-YFP(+) biofilms (see supplemental Figure S3C for calculation details). While the percentage of total CD43 area overlapping with Gag-YFP(+) signals was negligible (around 2%), 50% of the total area occupied by CD9 or CD81 colocalized with Gag-YFP(+) biofilms (Figure 3F). A milder but still significant colocalization was observed for CD82, with approximately 30% of its total area polarized in Gag-YFP(+) biofilms. Altogether, these results highlight that Gag expression is sufficient to initiate tetraspanins polarization and that CD9, CD81, and CD82 concentrate in HTLV-1 biofilms at the surface of infected T-cells.

### CD9, CD81, and CD82 are incorporated into individual HTLV-1 particles

As we showed that tetraspanins were released within HTLV-1 biofilms and clustered in their vicinity, we now examined if CD9, CD81, and CD82 were incorporated into the virus envelope. To this aim, we used STED microscopy and immuno-electron microscopy (IEM) which provide super-resolution imaging to visualize and discriminate individual virions. As we showed in Figure 1E using isolated biofilms, the diameter of individual Gag-YFP(+) virions measured by STED microscopy is 181nm ± 23nm.

First, C91-PL cells were stained for cortical F-actin, which highlights cell membrane boundaries, and for HTLV-1 Env to identify virions (Figure 4A). Viral biofilms appeared as dense structures enriched in mature virions (Env+) that were found at the surface of infected cells without being necessarily associated with the host cell plasma membrane (see supplemental Figure S4A for more acquisitions). This was confirmed by transmission electron microscopy imaging of HTLV-1 infected T-cells (Figure 4A) where viral biofilms concentrate virions and ECM components on top of the plasma membrane (see supplemental Figure S4B for more acquisitions). To test whether tetraspanins were similarly detected inside released particles accumulated in biofilms, C91-PL cells were electroporated with Gag-YFP and stained for CD9, CD81, CD82, or CD43 (Figure 4B). As opposed to CD43 which was only found at the plasma membrane and seemed to be excluded from viral aggregates, individual HTLV-1 particles were embedded in dense networks of CD9, CD81, and CD82 (magnified images in Figure 4B). This suggests that in addition to the plasma membrane, CD9, CD81, and CD82 also accumulate in the periphery of HTLV-1 particles. However, we could not exclude at this point that tetraspanins could be trapped in membrane protrusions within biofilms and not directly incorporated into HTLV-1 particles. To discriminate between these two hypotheses, Gag-YFP(+) biofilms were isolated; stained for Env, CD9, CD81, or CD82; and imaged by STED (Figure 4C). The three tetraspanins were observed at the periphery of isolated Gag-YFP(+) particles in an Env-like pattern, without being detected in the particles’ core (positive for Gag-YFP but negative for CD9, CD81, or CD82, Figure 4C and 4D). Moreover, we measured an equivalent distance between Env-Gag signals and Gag-tetraspanin signals using peak-to-peak distances analysis on cross-sections from Figure 4D. On average: Env-Gag distance = 114nm ±□39nm; CD9-Gag distance = 119nm ± 56nm; CD81-Gag distance = 112nm ± 45nm; and CD82-Gag distance = 105nm ± 33nm (Figure 4E). These results indicate that Env, CD9, CD81, and CD82 are equidistant from the particles’ center, suggesting that tetraspanins are incorporated into the viral envelope. This was confirmed by IEM (Figure 4F) where Env, CD9, CD81, and CD82 were all detected at the periphery of HTLV-1 particles contained in C91-PL cells biofilms (see supplemental Figure S4C for larger magnifications). While tetraspanins accumulated in HTLV-1 virions, we couldn’t detect any signal for CD9, CD81, or CD82 at the plasma membrane by IEM suggesting a specific enrichment of these molecules in viral particles (Figure 4F). In addition, the number of gold nanobeads per particle, which is relative to the number of target molecules per virion (Figure 4G), was around 1 gold-beads (± 1,2) per particle for CD9 and CD81 stainings, in comparison to a mean of 4 gold beads (±□ 2,5) per particle for Env and CD82 stainings (Figure 4G). Altogether, our findings highlight the Env-like incorporation of CD9, CD81, and CD82 into viral particles aggregated within isolated or cell-surface HTLV-1 biofilms.

**Figure 4:**
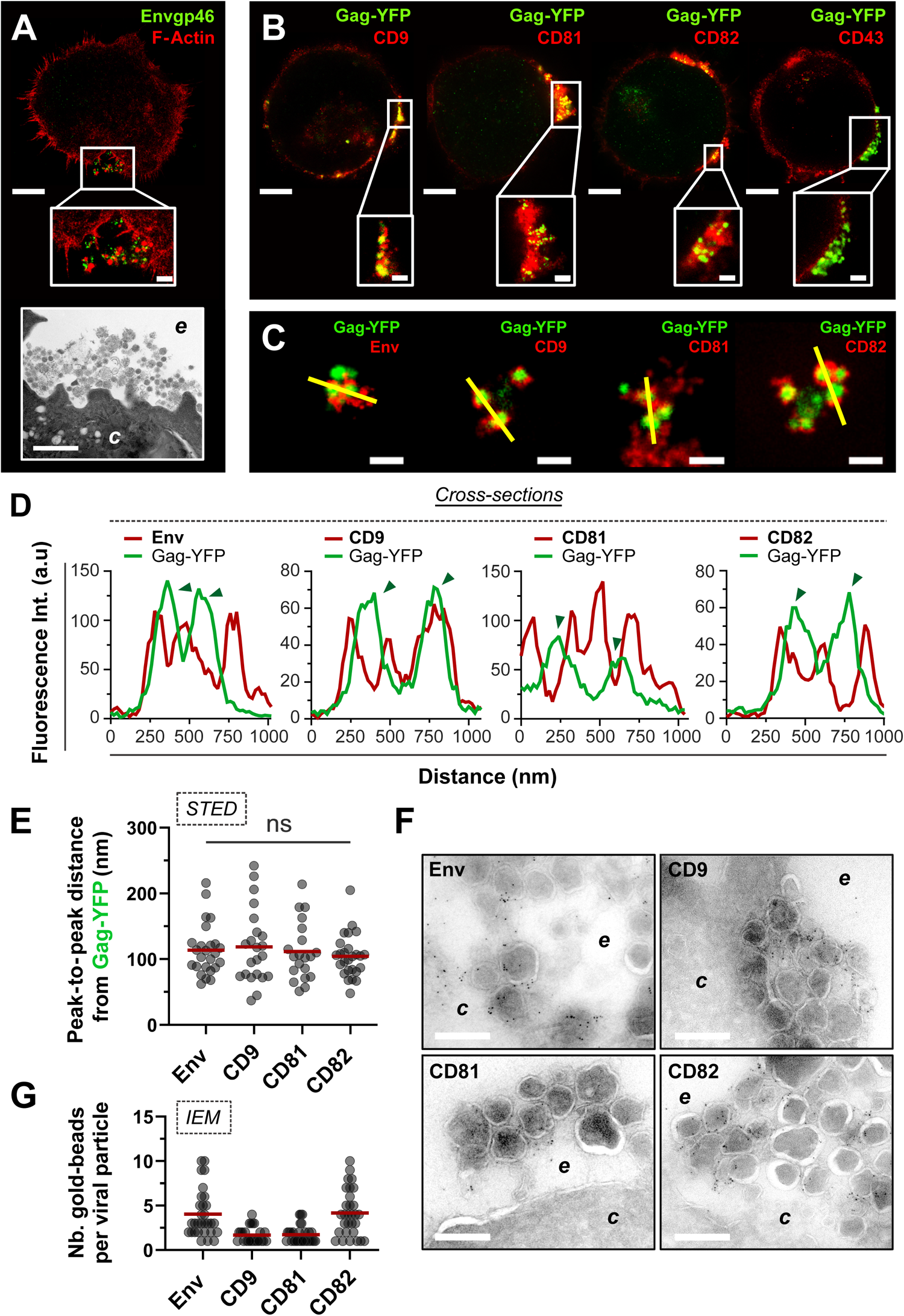
CD9, CD81, and CD82 are incorporated into individual HTLV-1 particles. **(A)** Top: Representative STED 2D image of an HTLV-1 infected T-cell (C91-PL) permeabilized and stained for cortical F-actin (red) and for HTLV-1 viral envelope (green). Scale bar = 5µm. Middle: Higher magnification of the Env(+) viral biofilm. Scale bar = 500nm. Bottom: Transmission electron microscopy image of HTLV-1 biofilm at the surface of a C91-PL cell. Scale bar = 1µm. *c* = cytosol; *e* = extracellular environment. **(B)** Top: Representative STED 2D images showing Gag-YFP(+) viruses embedded in CD9, CD81, or CD82 enriched domains at the surface of C91-PL cells. Scale bars = 5µm. Bottom: Zoomed images of viral biofilms. Scale bars = 500nm. **(C)** Representative STED 2D images of Gag-YFP(+) cell-free biofilms (isolated as detailed in Figure 1C) stained for Envgp46, CD9, CD81, or CD82 (red). Scale bars = 500nm. **(D)** Graphs corresponding to the yellow cross-sections in panel C. Arrows indicate individual particles (FWHM ≈ 150-200nm). **(E)** Scatter plot showing the distribution of the distances measured between the peak centers of Env/CD9/CD81/CD82 signals (extracted from panel C cross-sections) and the peak centers of Gag-YFP signals (n > 20 peak-to-peak distances measured for each condition). Mean = red bar. Kruskal-Wallis statistical tests were used to assess any significant difference: ns = non-significant. **(F)** Immuno-electron microscopy images showing viral biofilms at the surface of C91-PL cells stained for Env, CD9, CD81, or CD82 with antibodies coupled to 6nm gold beads. *c* = cytosol; *e* = extracellular environment. Scale bars = 200nm. FWHM: full width at half maximum. **(G)** Scatter plot showing the number of gold beads per viral particle (relative to the number of target molecules per virion). n > 25 particles per condition. Mean = red bar.

### CD82 regulates the polarization of HTLV-1 biofilms at the surface of infected T-cells

After having established the incorporation of CD9, CD81, and CD82 into viral particles within biofilms, we examined their potent functions in the organization of HTLV-1 biofilms at the cell surface, as tetraspanins are known to be molecular scaffolds that distribute partner proteins into highly organized microdomains (for a review see^43,44^). To this aim, we designed shRNAs delivered using GFP(+) lentivectors targeting each tetraspanin of interest to silence their expression in chronically infected T-cells (C91-PL). Non-transduced cells (wild-type, WT) along with cells transduced with scrambled shRNAs (shCtrl) were used as negative controls. The efficiency of shRNA-silencing in the cell population was controlled by western blot at day 5 post-transduction and reached repeatedly around 40% for shCD9, 70% for shCD81, and 50% for shCD82 (Figure 5A). Simultaneously, C91-PL cells were fixed and surface-stained for Env at day 5 post-transduction (Figure 5B). Silenced cells were identified by GFP expression, which was absent in WT C91-PL cells as expected (shown in representative images of Figure 5B). CD9 and CD81 silencing did not seem to affect the integrity of viral biofilms which remained large, polarized Env(+) aggregates similar to those observed in control cells (Figure 5B). However, CD82 silencing led to a complete reorganization of HTLV-1 biofilms, with smaller Env clusters scattered all around the cell surface (Figure 5B). This clear redistribution was also observed when HTLV-1 biofilms were stained for Gag in CD82-silenced C91-PL cells (Figure 5H).

**Figure 5:**
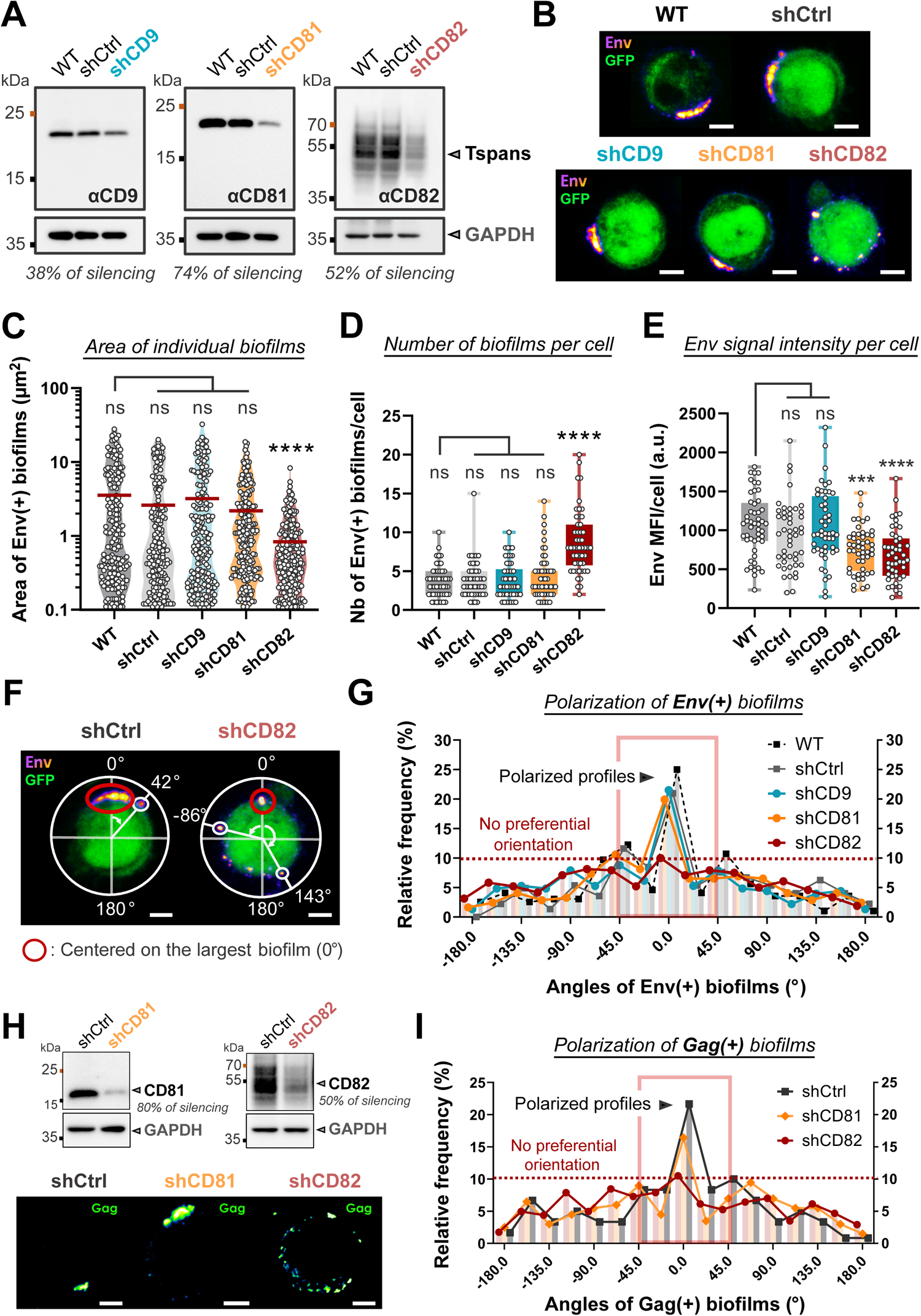
CD82 silencing induces the depolarization of HTLV-1 biofilms at the surface of infected T-cells. **(A)** Immunoblots showing CD9, CD81, and CD82 expression in C91-PL cells transduced or not (WT) with scrambled shRNAs (shCtrl) or shRNAs targeting CD9, CD81, or CD82, 5 days post-transduction. Endogenous GAPDH is used as a loading control. Silencing efficiency is reported below each immunoblot and was calculated as the ratio between the intensity of tetraspanins signals upon silencing and their expression in WT conditions (both normalized to GAPDH). **(B)** Representative confocal images of C91-PL transduced or not (WT, GFP-) with different shRNAs (GFP+) - shCtrl, shCD9, shCD81, or shCD82. All cells were fixed and stained for HTLV-1 Env (fire red) 5 days after shRNAs transduction. Scale bars = 5µm. **(C)** Violin plot showing the area (µm^2^) of individual Env(+) biofilms in control cells versus CD9/CD81/CD82 silenced cells (n > 180 biofilms per condition). Mean = red bar. **(D)** Box plot showing the number of Env(+) biofilms at the surface of control cells versus CD9/CD81/CD82 silenced cells (n > 50 cells in each condition). **(E)** Box plot showing the mean fluorescence intensity (MFI) of Env signal per cell in control cells versus CD9/CD81/CD82 silenced cells (n > 50 cells). **(F)** Representative confocal images showing examples of the analysis performed to assess the angular distribution of HTLV-1 biofilms at the cell surface (see “Confocal images analysis” in the Material & Methods section). **(G)** Frequency distribution of the angles of Env(+) biofilms at the surface of control cells (WT, shCtrl) versus CD9/CD81/CD82 silenced cells (n > 180 biofilms for n > 50 cells per condition). While a peak between −45° and 45° (red square) is representative of a polarized profile, the absence of a peak indicates a depolarized pattern (red dotted line). **(H)** Top: Immunoblots showing CD81 and CD82 expression in C91-PL cells transduced with scrambled shRNAs (shCtrl) or shRNAs targeting CD81 or CD82, 5 days post-transduction. Endogenous GAPDH is used as a loading control. Bottom: Representative confocal images of C91-PL transduced with different shRNAs - shCtrl, shCD81, or shCD82. All cells were fixed and stained for HTLV-1 Gag (green fire blue) 5 days after shRNAs transduction. Scale bars = 5µm. **(I)** Frequency distribution of the angles of Gag(+) biofilms at the surface of control cells (shCtrl) versus CD81/CD82 silenced cells (n > 120 biofilms for n > 30 cells per condition). Kruskal-Wallis statistical tests were used to assess any significant difference: ns = p.value > 0,05; *** = p.value ≤ 0.001; **** = p.value ≤ 0.0001.

Accordingly, CD82 silencing resulted in a significant decrease in the area of individual Env(+) biofilms (mean area = 1.1 µm^2^), as opposed to CD9 or CD81 silencing (mean area = 2.90µm^2^, Figure 5C). This was combined with a significant increase in the number of Env(+) biofilms per cell, from a mean of 4 biofilms/cell in controls to a mean of 9 biofilms/cell in CD82-silenced cells (Figure 5D). In addition, Env mean fluorescence intensity (MFI) measured at the cell surface of non-permeabilized C91-PL cells showed a 30% decrease upon CD81 and CD82 silencing as compared to the other conditions (Figure 5E). This suggests that CD81 and CD82 silencing might partially impair Env recruitment to the plasma and/or virions accumulation at the cell surface.

Next, we examined the distribution profiles of Env(+) biofilms at the surface of infected T-cells (C91-PL) to quantify HTLV-1 biofilm reorganization upon CD9, CD81, or CD82 silencing as exemplified in Figure 5F (see “Confocal images analysis” in the materials and methods section for analysis details). For WT, shCtrl, shCD9, and shCD81 conditions, around 45% of all Env(+) bio-films were detected close to 0° (which is positioned on the largest biofilm), indicating a preferential orientation at the cell surface (“polarized profile”, Figure 5G). However, upon CD82 silencing, Env(+) biofilms were redistributed at the cell surface with an increased number of viral aggregates further away from 0° and less than 30% between −45° and 45°, which indicates a depolarized pattern (“no preferential orientation”, Figure 5G). With a similar behavior observed for Gag clusters upon CD82 silencing (Figure 5I), we examined if endogenous CD82 was physically interacting with HTLV-1 structural proteins by performing coimmunoprecipitation assays using chronically infected T-cells (supplemental Figure S5). Although we did not detect any interaction between Gag and CD9 or CD81, our results show that HTLV-1 Gag repeatedly coimmunoprecipitated with CD82, which demonstrates their ability to form complexes in infected T-cells (supplemental Figure S5). Overall, our results demonstrate that tetraspanins play a critical role in the accumulation of virions at the cell surface and that CD82 is required to maintain HTLV-1 biofilm polarization, most probably through interactions with Gag.

### CD82 is required for efficient HTLV-1 cell-to-cell transmission

Biofilm polarization is hypothesized to be an important parameter for the bulk delivery of HTLV-1 particles through cell-to-cell contacts. To test this hypothesis, the efficiency of viral transmission was assessed by putting reporter Jurkat T-cells in co-culture with C91-PL cells silenced or not for the expression of CD81 or CD82 (Figure 6A). Reporter Jurkat T-cells (JKT-LTR-Luc) encode luciferase under the control of HTLV-1 long terminal repeat (LTR) promoter which is transactivated by the viral protein Tax during a productive infection^24^. Hence, luciferase activity is relative to the efficiency of viral transmission. Of note, CD9 was excluded from the analysis at this point, as its silencing did not impact viral biofilm organization or HTLV-1 Env expression at the cell surface (Figure 5B and 5G). Luciferase activity was measured after 24 hours of co-culture between reporter cells and CD81- or CD82-silenced C91-PL cells at days 3, 5, and 7 post-transductions. The efficiency of CD81 and CD82 silencing was controlled by western blot and reached up to 70% as compared to their expression in WT cells (Figure 6B).

**Figure 6:**
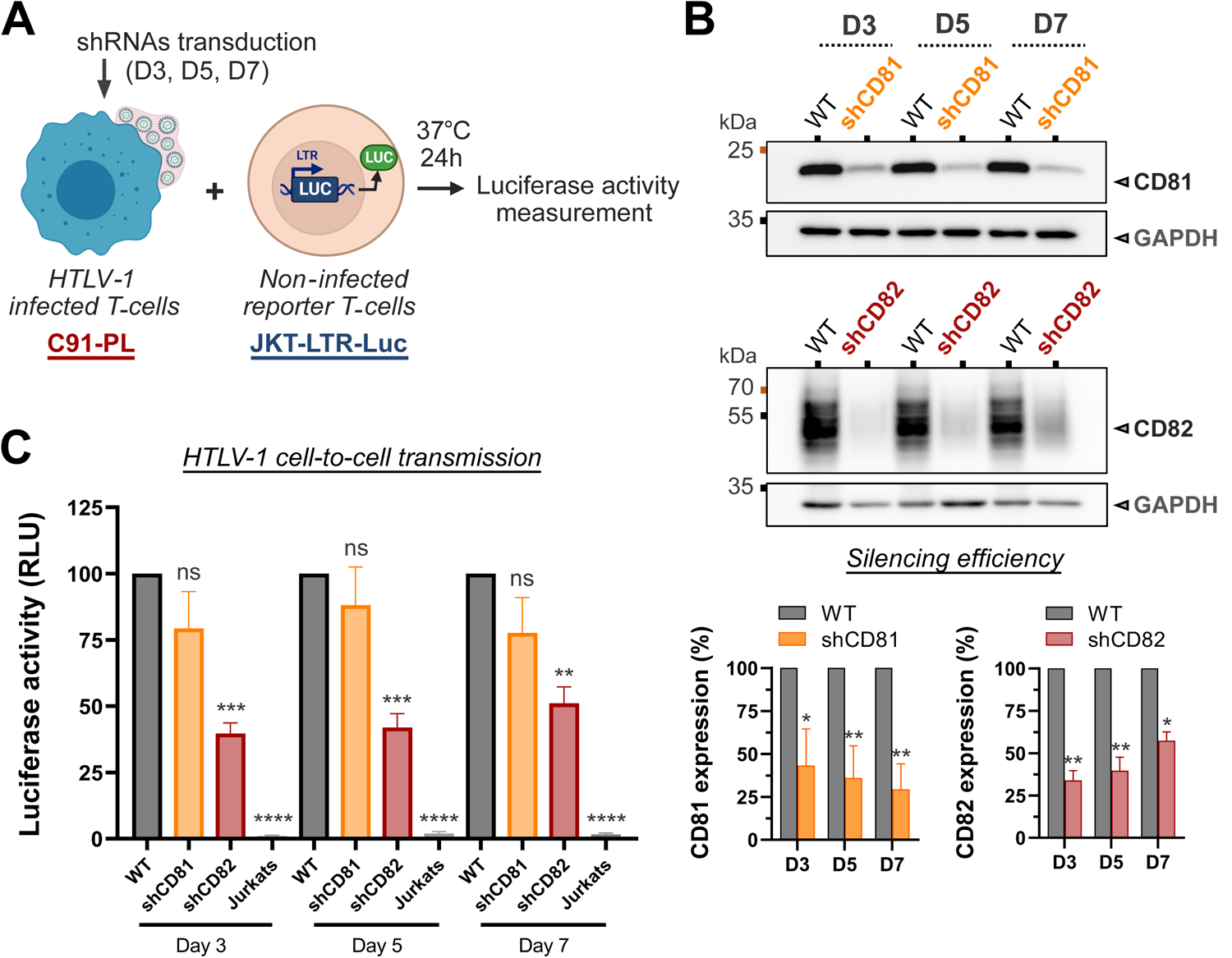
CD82 is required for efficient HTLV-1 cell-to-cell transmission. **(A)** Luciferase reporter assay scheme: C91-PL cells transduced or not with shRNAs targeting CD81 or CD82 for 3, 5, or 7 days, were co-cultured with non-infected reporter Jurkat T-cells for 24h. Reporter cells are stably transfected with a plasmid bearing the luciferase gene downstream of the HTLV-1 LTR promoter which is transactivated upon infection by the viral transactivator protein Tax. **(B)** Western blots showing CD81 or CD82 expression in shRNA-transduced C91-PL cells versus control cells (WT: non-transduced cells). Endogenous GAPDH is used as a loading control. The mean expression ± SEM of CD81 and CD82 after shRNA-silencing at days 3, 5, and 7 post-transductions are reported on the graphs below (n = 4 independent experiments). **(C)** Luciferase activity of HTLV-1-LTR Luc reporter T-cells co-cultured with WT C91-PL cells, shRNAs-transduced C91-PL cells, or WT Jurkat T-cells, as a measure of HTLV-1 cell-to-cell transmission efficiency. Results were obtained from four independent experiments performed in triplicates with different lentivector productions. The histogram shows the mean luciferase activity ± SEM for each condition in %, normalized to the positive control (JKT-LTR-Luc co-cultured with WT C91-PL cells). Ordinary one-way ANOVA statistical tests were used to assess any significant difference: ns = p.value > 0,05; * = p.value ≤ 0.05; ** = p.value ≤ 0.01; *** = p.value ≤ 0.001; **** = p.value ≤ 0.0001.

For each time point, we observed no significant difference in the luciferase activity of reporter cells after co-culture with WT C91-PL cells or CD81-silenced C91-PL cells (Figure 6C). In contrast, luciferase activity was severely reduced upon co-culture with CD82-silenced C91-PL cells (Figure 6C). Importantly, the infectivity of C91-PL cells correlated with CD82 silencing efficiency, demonstrating the direct impact of CD82 expression on HTLV-1 cell-to-cell transmission. For instance, the 67% (□±□6%) decrease in CD82 expression at day 3 post-transduction resulted in a 61% (□±□4%) reduction of HTLV-1 transmission, whereas a less efficient CD82 knockdown at day 7 post-transduction (around 43% ±□5%) led to a 49% (□±□6%) reduction in infectivity (Figure 6C). To control that viral budding was not differentially impacted by the loss of CD81 or CD82 expression, we quantified HTLV-1 Gag release by western blot upon CD81 and CD82 silencing (supplemental Figure S6A). A small decrease in the viral release was noticed in both conditions as compared to the control (around 20%, supplemental Figure S6B). This relative decrease was mainly due to an increased proportion of Gag retained in the cell lysate (supplemental Figure S6C), supporting the hypothesis that CD81 and CD82 silencing might partially inhibit virions accumulation at the cell surface and release, which is consistent with the observations made in Figure 5E. Nevertheless, the significant reduction in luciferase activity measured strictly upon CD82 silencing resulted from impaired cell-to-cell infectivity, which indicates a crucial role for CD82 tetraspanin in HTLV-1 intercellular transmission.

## DISCUSSION

Polarized aggregates of infectious HTLV-1 virions, identified as biofilms, have been observed in HTLV-1 infected T-cell lines and at the surface of primary CD4(+) T-cells isolated from HAM/TSP patients or asymptomatic HTLV-1 carriers^24^, suggesting a conserved role in HTLV-1 pathogenesis. Besides this pioneering discovery, the molecular and functional description of viral biofilms is still in its infancy, though of utmost importance to fully understand retroviruses’ pathogenesis. In this study, we explored the molecular composition of viral biofilms and identified specific cellular components (*i.e.,* tetraspanins) that control the cell-surface distribution and the intercellular transmission of HTLV-1 biofilms.

Here, we first demonstrated that CD9, CD81, and CD82 tetraspanins were enriched in HTLV-1 biofilms (Figure 2) and clustered in their vicinity at the cell surface (Figure 3). Interestingly, tetraspanin-enriched microdomains (TEMs) pre-exist in the plasma membrane of lymphocytes, where more than three different molecules of this family can reside in a single complex^45^. It is noteworthy that several viruses budding from the plasma membrane like the herpes simplex virus 1 (HSV-1)^46^, the influenza A virus (IAV)^47^, and HIV-1^30,48^ assemble in TEMs. Therefore, TEMs containing CD9, CD81, and CD82 could be preferentially targeted by HTLV-1 structural proteins and act as exit platforms. Accordingly, earlier reports showed that both HTLV-1 Gag and Env proteins could interact with tetraspanins^33,34^. For instance, the HTLV-1 Gag matrix domain was reported to interact with the intracellular loops of both CD81 and CD82 when transiently over-expressed in HEK293T cells^33^. Here, we showed that ectopic Gag-YFP expression in HTLV-1 infected T-cells deficient for viral structural protein expression was sufficient to relocate CD81 and CD82 within Gag clusters at the cell plasma membrane (Figure 3). This suggested that Gag could also interact with CD81 and CD82 in infected T-cells. However, our coimmunoprecipitation experiments in HTLV-1 infected T-cells only confirmed the interaction of Gag with CD82, suggesting that Gag might have more affinity for endogenous CD82 than for CD81 in infected T-cells (Figure S5). Consequently, CD81 recruitment to Gag clusters might be bridged by CD82. In addition, CD82 could be recruited by HTLV-1 Env, as it was reported that they both associate early along CD82 biosynthesis pathway up to the cell plasma membrane^34^. This interaction was proposed to account for HTLV-1 Env localization and concentration in TEMs. Here, we propose that CD82 might form a complex with both Gag and Env (Figures 5 and S5), with Gag being a crucial determinant for the initiation of CD82 polarization (Figure 3). Importantly, other viruses targeting TEMs recruit different tetraspanins. For instance, IAV specifically recruits CD81 to its assembly sites^47^, and HIV-1 Gag and Env accumulate in TEMs^31,48^ mainly through interactions with CD81^30^. Interestingly, while HIV-1 Gag could also interact with CD63 and CD82^30^, HIV-1 Env did not interact with CD82^34^, highlighting different but specific molecular determinants involved in viral proteins-tetraspanins interplay.

Interactions between viral proteins and tetraspanins within TEMs frequently translate into their incorporation into budding virions. This was previously reported for several enveloped viruses such as the hepatitis C virus (HCV), IAV, and HIV-1 (for a review see^49^). For example, in addition to CD81 which interacts with HIV-1 Gag and Env^30,48^, HIV-1 particles incorporate CD9, Tspan14, CD53, CD63, and CD82^50,51^. Similarly, we showed that CD9, CD81, and CD82 are incorporated into HTLV-1 particles (Figure 4) suggesting that interactions between HTLV-1 structural proteins and tetraspanins are likely to be maintained through the budding process. Moreover, both CD81 and CD82 seem to play a role in HTLV-1 budding process as their silencing induces a reduction in Env expression at the cell surface (Figure 5) along with a small decrease in viral release (Figure S6). Likewise, HIV-1 release is decreased by 20% upon CD81 silencing which demonstrates that these tetraspanins promote retroviruses’ egress^31,52^. Taken together, these observations suggest that tetraspanin webs may facilitate Env glycoproteins uptake, HTLV-1 assembly, and budding during virus morphogenesis through the induction of large assembly platforms that concentrate both tetraspanins and viral proteins.

In addition to their potential function in HTLV-1 assembly, we showed that tetraspanins could modulate viral clustering and HTLV-1 biofilm polarization at the cell surface (Figure 5). Examples of viruses that concentrate within TEMs in discrete plasma membrane regions were reported in the literature: for instance, IAV infection redistributes CD81 on the plasma membrane into concentrated patches of viral assembly and budding sites^47^. Also, HIV-1 Gag expression induces the concentration of CD9 and CD81 nanoclusters within Gag assembly areas^53^. Here, we observed that the transient expression of HTLV-1 Gag-YFP in infected T-cells that do not produce viral particles (nor biofilms) initiates CD81 and CD82 polarization within Gag clusters (with a similar but smaller effect for CD9). Taken together, this suggests that retroviral Gag proteins have the potential to drive tetraspanins clustering and polarization. Here, this phenomenon could be indirect for CD9/CD81, and direct for CD82 which can form complexes with HTLV-1 Gag (Figure S5). Importantly, while CD9 or CD81 silencing had no effect on HTLV-1 biofilm or-ganization at the cell surface, CD82 silencing induced a complete reorganization of viral aggregates (Figure 5). Similar effects have been reported for HIV-1 where CD81 silencing induces a redistribution of HIV-1 Gag clusters at the surface of infected T-cells^30^. Here, CD82, but not CD81, is required to maintain the architecture and polarization of HTLV-1 biofilm, highlighting distinct molecular events involved in HIV-1 and HTLV-1 clustering. This observation is consistent with the ability of Env and Gag viral proteins to interact preferentially with CD81 (for HIV-1) or CD82 for (HTLV-1). Consequently, the concomitant aggregation of CD82 and assembling Gag molecules into large platforms at the cell plasma membrane could promote HTLV-1 Gag-Env encounters leading to the production of dense biofilms containing mature particles. Comparable events were reported for HIV-1, where Gag assembly is known to induce the aggregation of small Env clusters into larger domains that are completely immobile^54^.

The molecular mechanism ruling the CD82-dependent polarization of HTLV-1 biofilms remains to be elucidated. CD82 can form multimeric complexes with other tetraspanins but also with integrins and heparan sulfate proteoglycans^55–57^, which are adhesion molecules also enriched in HTLV-1 biofilms^24^. Thus, tetraspanins, which modulate intercellular adhesion processes (especially CD82^58^) could direct virions assembly to biofilms preferentially formed at intercellular junctions (as shown in Supplemental Video S1, Supplemental Figure S1). Additionally, the differential effects observed for CD9, CD81, and CD82 could arise from their three-dimensional structure and subsequent dynamics in the cell plasma membrane. These three tetraspanins, which share conserved topologies, differ in terms of post-translational modifications in their large extracellular loop (LEL). For example, CD82 LEL is highly N-glycosylated as opposed to CD9 and CD81 LELs^59^, especially in HTLV-1 infected T-cells^34^ where the MGAT-3 glycosyltransferase responsible for CD82 N-glycosylation^56^ is up-regulated (GSE17718 database^40^). Interestingly, CD82 N-glycosylation is known to promote CD82-integrins interactions^56^, modulate CD82 membrane clustering^60^, and participate in tetraspanins complex formation^61^. Moreover, increased glycosylation of extracellular domains is directly linked to a decrease in the diffusivity of transmembrane proteins^62^. Accordingly, CD9 and CD81 dynamics are inferior to CD82 dynamics in the plasma membrane of HB2 cells^63^. Therefore, we can speculate that CD82 N-glycosylation could promote its interaction with membrane or submembrane components (*i.e.*, integrins) and restrict its dynamics at the cell surface. Both processes could thus lead to a subsequent decrease in the local diffusion of partner molecules, which could ultimately contribute to the pivotal role of CD82 in the maintenance of HTLV-1 biofilm architecture. Interestingly, after treating HTLV-1 infected T-cells with an inhibitor targeting the oligosaccharyltransferase (OST), which is responsible for initiating N-glycosylation of membrane proteins, we observed CD82 deglycosylation along with a complete depolarization of viral biofilms (Figure S7). This supports the notion that N-glycosylation of host cell membrane proteins, including CD82, is required for the polarization of HTLV-1 biofilms (Figure S7).

Finally, we demonstrated that the CD82-dependent clustering of viral particles within TEMs at the cell surface is a key parameter in HTLV-1 cell-to-cell transmission (Figure 6). Early electron microscopy studies revealed that other viruses could also form viral aggregates at the cell surface resembling viral biofilms, including IAV^64^, vesicular stomatitis virus^65^, and HIV-1^52,66^. Clusters of mature HIV-1 particles resembling biofilms were also detected at the surface of infected MOLT T-lymphoblastic cells^30^, at the surface of primary CD4(+) T-cells upon *in vitro* HIV-1 infection^66,67^, and at the surface of CD4(+) T-cells from HIV-1-infected patients^66^. Virions aggregation significantly increases the chances of multiple viral genomes being jointly delivered to target cells, as shown recently for VSV^68^ and HIV-1^66^, which demonstrates the ability of enveloped viruses to establish collective infectious units. Accordingly, viral biofilms account for 80% of HTLV-intercellular transmission *in vitro*^24^, and our results further support the importance of the collective transmission of clustered virions. Moreover, our results provide a molecular basis for this process, as CD82 silencing, which provokes HTLV-1 biofilm reorganization, severely reduces the ability of infected T-cells to transmit the virus (by up to 60%, Figure 6). In contrast, CD81 silencing, which does not affect HTLV-1 biofilm clustering (Figure 5), has no significant effect on HTLV-1 cell-to-cell transmission (Figure 6). Altogether, our results suggest that viral biofilm polarization is required for efficient cell-to-cell transmission. Different mechanisms which are not fully deciphered yet may account for the initiation and maintenance of virus polarization in different viral infections. For HTLV-1, our overall observations provide a specific role for CD82 in the regulation of HTLV-1 biofilm polarization and its subsequent intercellular transmission.

## MATERIAL & METHODS

### Cell lines

C91-PL cells (HTLV-1 chronically infected T-cell line, Cellosaurus, ref CVCL_0197), MT-2 cells (HTLV-1 chronically infected T-cell line^69^, NIH, ref 237), and C8166 cells (HTLV-1 infected T-cell line impaired for infectious particles’ production^70^, ECACC ref 88051601) were obtained from Helene Dutartre’s lab and cultured in 75 cm^2^ flasks in BSL3 facilities at 37°C with 5% CO_2_. Jurkat cells (non-infected T-lymphocytes, from ATCC, ref ACC 282) were also cultured in 75 cm^2^ flasks in BSL2/BSL3 facilities at 37°C with 5% CO_2_. Jurkat cells transfected with a plasmid encoding the luciferase enzyme (Luc) under the control of HTLV-1 long terminal repeat (LTR) promoter, which is transactivated by HTLV-1 Tax (Jurkat-LTR-Luc^24^), were grown under hygromycin selection (450 μg/mL, Invivogen). All T-lymphocytes cell lines were maintained in complete RPMI medium: RPMI GlutaMAX (ThermoFisher Scientific) supplemented with decomplemented FCS 10% (ThermoFisher Scientific) and 1% penicillin/streptomycin (Merck). Human embryonic kidney 293T cells (HEK 293T) were maintained in DMEM GlutaMAX (Dulbecco’s Modified Eagle Medium, ThermoFisher Scientific) supplemented with decomplemented FCS 10% (ThermoFisher Scientific) and 1% penicillin/streptomycin (Merck) at 37°C with 5% CO_2_.

### Plasmids and DNA constructs

The plasmid encoding HTLV-1 Gag-YFP under the CMV promoter (pHTLV Gag-YFP) was described previously^71^ and is a kind gift from Dr. David Derse’s lab. The pFG12 GFP(+) vectors encoding the control shRNAs and shRNAs targeting CD81 were a kind gift from Dr. Birke Bartosch’s group (Inserm U1052, Cancer Research Centre of Lyon, France). Plasmids encoding shRNAs targeting CD9 and CD82 were designed in the lab as follows: shCD9 and shCD82 synthetic sequences (designed and ordered on GeneArt, ThermoFisher Scientific) were placed downstream the U6 promoter and cloned between XbaI and HpaI restriction sites into the pFG12 GFP(+) lentiviral vector (obtained from Dr. David Baltimore’s lab, Addgene plasmid #14884). All constructs were verified by DNA sequencing (Eurofins genomics, Plasmidsaurus). The packaging vectors pVSV-G (encoding the vesicular stomatitis virus envelope glycoprotein) and p8.2 (encoding HIV-1 Gag-Pol and its accessory proteins except for Vpu) were used for vectorization (as it was previously described^72,73^) of the different shRNAs into lentiviral particles (see below).

### Antibodies

All antibodies used in this study along with their targets, species, clone numbers, references, initial concentrations, and working dilutions are listed in Supplemental Table S1.

### Electroporation of HTLV-1 chronically infected cells for microscopy

Electroporation was conducted using the Gene Pulser Xcell™Electroporation System (BioRad), according to the following protocol: cells were washed in 1X Phosphate buffered saline (PBS, ThermoFisher Scientific), counted, and resuspended in Opti-MEM (1X) Reduced Serum Medium (ThermoFisher Scientific) at a concentration of 2 x 10^7^ cells/mL. 8 x 10^6^ cells (*i.e.,* 400µL) were then mixed with 10µg plasmid for each condition. The resulting mixtures were put at 37°C for 30min and transferred to 0.4 cm electroporation cuvettes (Gene Pulser/MicroPulser, BioRad), which were submitted to electroporation using 10ms pulses at 180V repeated three times at one-second intervals. Cells were then gently transferred to 6-well plates in pre-warmed RPMI GlutaMAX (ThermoFisher Scientific) 10% FCS without P/S and cultured at 37 °C with 5% CO_2_. After 24-48 hours, cells were washed in 1X PBS and plated on 0.01% poly-L-lysine (Sigma-Aldrich) coated glass-bottom Fluorodishes (WPI) for 30min at room temperature. Samples were then fixed using 4% paraformaldehyde for 15min at room temperature.

### Living cells imaging

Living C91-PL cells electroporated with HTLV-1 Gag-YFP construct were plated in uncoated 6 wells-plates in complete RPMI medium 20 hours post-electroporation. Plates were placed in the incubator chamber of an automated laser-scanning confocal microscope (Cell-Discoverer 7 LSM900, Zeiss, Germany). Epifluorescence images were generated every 15min for 5 hours (supplemental Video S1 and supplemental Figure S1).

### Isolation of viral biofilms for mass spectrometry or microscopy

Viral biofilms were isolated from the supernatants of chronically infected cell lines (wild-type for mass spectrometry or electroporated with HTLV-1 Gag-YFP for microscopy). Cells were cultured in 25 cm^2^ flasks in RPMI complete medium at an initial concentration of 0.5 x 10^6^ cells/mL. Cultures were left untouched for 96h to favor biofilm enrichment and its release in the culture supernatant. Cells were removed from the culture by centrifugation at 1000g for 5min. Supernatants were collected and then submitted to a 10000g centrifugation for 40min. Pellets containing biofilms were resuspended and used immediately. For microscopy experiments, pellets were resuspended in TNE 1X (Tris/HCl pH7.4 NaCl, EDTA), plated on 0.01% poly-L-lysine (Sigma-Aldrich) coated glass-bottom Fluorodishes (WPI) for 30min at room temperature, and fixed using 4% paraformaldehyde for 15min. For mass spectrometry experiments, 10000g pellets containing biofilms were resuspended in 1X PBS and inactivated at 56°C for 30min.

### Atomic force microscopy coupled with fluorescence imaging

Biofilms isolated from Gag-YFP(+) expressing C91-PL cells were diluted in TNE 1X (Tris/HCl pH7.4 NaCl, EDTA) and plated on glass-bottom fluorodishes (WPI) coated with 0.01% poly-L-lysine (Sigma-Aldrich). AFM imaging was performed on a NanoWizard IV atomic force microscope (Bruker) at room temperature. Fluorescence imaging was performed by wide-field illumination using a Nikon Ti-U inverted microscope mounted on the AFM machine and equipped with a 100X, 1.4 NA oil objective (Nikon). The AFM tip position was calibrated with the optical image using a specific software module (DirectOverlay, JPK Bio-AFM, Bruker). AFM topograph-ic images were acquired in quantitative imaging (QI) mode using BL-AC40TS cantilevers (Olympus). Cantilevers’ sensitivity and spring constant (kcant□=□0.1 N/m) were calibrated using the thermal noise method. The force applied to the samples was kept at 200 pN with an approach/retract speed of 25 µm/s (z-range = 100 nm). For image analysis, we used the JPK-data processing software (Bruker). Images were flattened with a histogram line fit and minor noise was removed using low-pass Gaussian and median filtering. Z-color scales (Figure 1D) are shown as relative after processing. To measure the maximal central height of each particle we used the cross-section tool of the analysis software.

### Mass spectrometry experiments

Biofilms isolated from C91-PL cells were diluted in buffer A (0.1% formic acid). Samples were injected for in-line analysis using nano-flow high-performance liquid chromatography (RSLC U3000, ThermoFisher Scientific) coupled to a mass spectrometer equipped with a nanoelectrospray source (Q Exactive HF, ThermoFisher Scientific). Peptides were separated on a capillary column (0.075 mm × 500 mm, Acclaim Pepmap 100, reverse phase C18, NanoViper, ThermoFisher Scientific) according to a 2-40% buffer B gradient (0.1 % formic acid, 80% acetonitrile) at a flow rate of 300 nL/min during 128 min. Spectra were recorded using the Xcalibur 4.1 software (ThermoFisher Scientific) with the 128LowQtt.meth method. Spectral data of the three independent experiments were analyzed using MaxQuant v2030 and Perseus v161043, with the use of the leading FPP v3.4 script. As a reference database, we used: RefProteome_HUMAN-cano_2022_01_UP000005640.fasta and UniProt-taxonomy 11908_202111.fasta (Human T-cell leukemia virus type 1) with the following fixed modification: Carbamidomethylation (C) and the following variable modifications: Acetyl (Protein N-term); Oxidation (M). Proteins identified with at least 2 peptides were kept. Also, proteins matching with a “Contaminants” database (keratin, trypsin, etc.) were removed from the analysis. The whole experimental process was conducted by Serge Urbach and Mathilde Decourcelle (Montpellier FPP platform).

### Mass spectrometry results analysis

Raw mass spectrometry data were filtered to remove “contaminants” including mitochondrial and ribosomal proteins, and the 100 most abundant proteins were selected based on their IBAQ score (which is the intensity averaged by the number of peptides that can be detected for a given protein). To only keep candidates detected with high confidence, molecules identified with less than three peptides were excluded from the analysis. The remaining 69 candidates were then matched against a list of proteins known to be up-regulated by HTLV-1 infection (GSE17718 database, Kress et al, 2009) and compared to potent HTLV-1 biofilm components identified in the literature.

### shRNAs-carrying lentivectors production and transduction

3×10^6^ HEK 293T cells were plated in 10cm culture dishes and cultivated overnight. Cells were then co-transfected using CaCl_2_/HBS 2X with plasmids (see “Plasmids and DNA constructs”) encoding VSV-G envelope (pVSV-G), HIV-1 Gag-Pol (p8.2), scrambled shRNAs, or shRNAs directed against the following targets: CD9, CD81, or CD82 (shRNAs sequences are listed in Supplemental Table S1). Culture media containing lentiviral particles were collected 48h posttransfection and filtered through 0.45 μm membranes. Particles were then purified by ultracentrifugation at 100000g for 1h30 at 4°C on a 25% sucrose cushion. Pellets were resuspended in 1X TNE (Tris/HCl pH7.4 NaCl, EDTA). The concentration of particles was determined by Videodrop (Myriade). C91-PL cells were transduced with the shRNAs-carrying lentivectors using a total of 100 particles per cell. 24h post-transduction cells were washed and cultured in RPMI complete medium for 72h (D3), 120h (D5), and 168h (D7). Then, cultures were centrifuged at 1000g for 5min. Cells were collected, washed, and resuspended in 1X PBS at a concentration of 1 million cells/mL. 500µL of cells were plated on 0.01% poly-L-lysine (Sigma-Aldrich) coated glass-bottom Fluorodishes (WPI) for immunofluorescence or resuspended in RIPA lysis buffer for western blots (see below). Supernatants were collected and centrifuged at 10000g for 40min to pellet biofilms. Biofilms-containing pellets were resuspended in 1X PBS and plated on glass-bottom fluorodishes (WPI) for immunofluorescence or resuspended in PBS 0.2% Triton for western blots (see below).

### Immunofluorescence for confocal microscopy

Fixed C91-PL cells (WT, transduced with shRNAs, or electroporated with Gag-YFP) plated on fluorodishes were incubated in 50mM NH_4_Cl for 5min at room temperature (RT) to quench free aldehydes and in 1X PBS containing 3% Bovine Serum Albumin (BSA, Sigma-Aldrich) for 15min at RT to block non-specific epitopes. After washing with PBS 3% BSA, samples were stained using 1µg/mL of primary mouse antibodies (αEnvgp46, αCD9, αCD81, or αCD82; see supplemental Table S1 for references) or rabbit antibodies (αCD43) diluted in PBS 3% BSA for 1 hour at RT. After several washes with PBS 3% BSA, samples were incubated with 1µg/mL of secondary antibodies (AlexaFluor647 anti-mouse, ThermoFisher, or Rabbit-Atto647N antirabbit, Sigma-Aldrich) for 1 hour at RT. Finally, samples were washed with 1X PBS and stored at 4°C in 2mL PBS until acquisition. Confocal images were obtained using a laser-scanning confocal microscope (LSM980, Zeiss) equipped with a 63X, 1.4 NA oil objective. All images were processed using ImageJ software (Fiji).

### Confocal images analysis

For colocalization analyses in Figure 3F, a schematic representation detailing the segmentation process and the calculation of the area of intersection between the two channels is provided in Supplemental Figure S3C. To analyze Env clusters at the surface of HTLV-1 infected cells transduced with scrambled shRNAs or shCD9/shCD81/shCD82 in Figure 5, we used ImageJ software (Fiji). Background noise was excluded using a mask in which only intensities above (Min + 0.2*(Max-Min)) were kept (Min: minimum pixel intensity, Max: maximum pixel intensity). After thresholding, the area of each cluster was extracted using the “Analyze particles” command on ImageJ (Figure 5C). The number of viral clusters per image (and thus per cell) after thresholding was used to quantify the number of Env clusters/cell (Figure 5D). The (total) MFI of the Env signal per image (and thus per cell) was extracted using ImageJ without thresholding (Figure 5E). For polarization analyses, an automated macro (considering the cell as a circle) attributed an angle between −180° and 180 to each Env cluster encountered on the cell surface with respect to the largest one positioned at 0° (representative images in Figure 5F). The frequency distribution of all angles (Figure 5G) was then obtained using Prism software (GraphPad).

### Immunofluorescence for 2D STED super-resolution microscopy

Fixed isolated biofilms and/or whole C91-PL cells (electroporated with Gag-YFP) plated on 0.01% poly-L-lysine (Sigma-Aldrich) coated fluorodishes were incubated in 50mM NH_4_Cl for 5min at room temperature (RT) to quench free aldehydes; incubated in PBS 3% BSA supplemented with 0.05% Saponin for 15min at RT to block non-specific epitopes and permeabilize the cells; and washed once using the same buffer. Then, cells were stained for 1h30 at RT with primary mouse antibodies (αEnvgp46, αCD9, αCD81, or αCD82), rabbit antibodies (αYFP), or actin-staining dyes (STAR red phalloidin, Abberior) diluted in PBS 3% BSA 0.05% Saponin. After three washes with PBS 0.05% Saponin 3% BSA, samples were incubated with the appropriate secondary antibodies (STAR Orange anti-rabbit, STAR Orange anti-mouse, or STAR Red anti-mouse, Abberior) diluted in PBS 3% BSA 0.05% Saponin for 1h30 at RT in the dark. Finally, samples were washed three times with 1X PBS and stored at 4°C in 2mL PBS until acquisition. Dual-color STED 2D images were acquired on a STED super-resolution microscope (Abberior Instruments Expert Line GmbH) equipped with a 100X oil objective using 580nm (Star-Orange) and 630nm (Star-Red) excitation laser sources, coupled with a pulsed 775nm STED laser. Using 25% of STED laser power, we could obtain a lateral resolution of less than 100nm. All images were processed with ImageJ software (Fiji).

### Transmission electron microscopy

HTLV-1 chronically infected T-cells (C91-PL) were fixed using 1% glutaraldehyde, 4% paraformaldehyde (Sigma-Aldrich), in 0.1 M phosphate buffer (pH 7.2) for 24h. Then, samples were washed in phosphate-buffered saline (PBS) and post-fixed using 2% osmium tetroxide (Agar Scientific) for 1h. Cells were then fully dehydrated in a graded series of ethanol solutions and propylene oxide. They were impregnated with a mixture of (1∶1) propylene oxide/Epon resin (Sigma-Aldrich) and left overnight in pure resin. Samples were then embedded in Epon resin (Sigma-Aldrich), which was allowed to polymerize for 48h at 60°C. Ultra-thin sections (90 nm) of these blocks were obtained with a Leica EM UC7 ultramicrotome (Wetzlar, Germany). Sections were stained with 2% uranyl acetate (Agar Scientific), 5% lead citrate (Sigma-Aldrich), and observed under a transmission electron microscope (JEOL 1011).

### Immuno-electron microscopy assay

HTLV-1 chronically infected T-cells (C91-PL) were fixed using 4% paraformaldehyde (PFA) in 1X PBS for 2h, washed twice with 1X PBS, and centrifuged for 10 min at 300g. Cell pellets were embedded in 12% gelatin and infiltrated overnight with 2.3 M sucrose at 4°C. Ultra-thin cryosections (90 nm) were obtained using a LEICA FC7 cryo-ultramicrotome at□−□110 °C. Sections were retrieved in a mix of 2% methylcellulose supplemented with 2.3 M sucrose (1:1) and deposited onto formvar-/carbon-coated nickel grids. Sections were incubated at 37°C to remove gelatin and labeled with primary antibodies targeting HTLV-1 Env, CD9, CD81, or CD82. Then, grids were washed with 1X PBS and labeled with secondary antibodies coupled to 6nm-diameter gold nanoparticles. Grids were washed using 1X PBS, post-fixed with 1% glutaraldehyde, and rinsed in distilled water. Contrast staining was achieved by incubating the grids with a mix of 2% uranyl acetate supplemented with 2% methylcellulose (1:10). Treated sections were then imaged under a transmission electron microscope (JEOL 1011).

### Western blot analysis

Before immunoblotting, HTLV-1 chronically infected cells (C91-PL) were washed in 1X PBS, pelleted at 1000g for 5min, and lysed in RIPA buffer (Sigma-Aldrich). The total protein concentration in cell lysates was measured using a Bradford protein assay kit (ThermoFisher). Biofilm fractions isolated from cell culture supernatants were resuspended in PBS 0.2% Triton (Sigma-Aldrich). Proteins from cell lysates (20µg per condition) or biofilm fractions (identical volume in each condition) were loaded on 10% acrylamide gels and separated by SDS-PAGE. Separated proteins were transferred onto polyvinylidene difluoride membranes (ThermoFisher Scientific) using wet transfer with tris-glycine-methanol buffer (milli-Q H_2_O supplemented with 15% methanol and 10% 10X Tris-glycine solution, Euromedex). Then, membranes were incubated for 30min in Tris-Buffered Saline-Tween (TBS-Tween) solution (milli-Q H_2_O 10% Trizma base-HCl 1M pH8 supplemented with 0.1% Tween 20, Sigma-Aldrich) supplemented with 5% milk to saturate non-specific sites and incubated with the corresponding primary antibodies (αGagp19, αCD9, αCD81, or αCD82) overnight at 4°C. Blotting membranes with antibodies targeting endogenous GAPDH was used as a loading control for cell lysates. After several washes in 5% milk TBS-Tween, membranes were incubated for 2h at room temperature with Horse Radish Peroxidase (HRP)-conjugated αMouse antibodies (Dako). After washing with TBS-Tween, HRP activity was revealed using ECL Prime reagents (Amersham). Images were acquired on a Chemidoc Imaging system (BioRad). Each band intensity was measured using the ImageJ software. Quantification of viral release was established by comparing the intensity of the Gagp19 signal detected in biofilms to the Gagp19 signal in both fractions (CL: Cell Lysates or BF: Biofilms) with the following formula: *(Gagp19 BF / (Gagp19 BF + Gagp19 CL))*100*. Each time point was normalized to its corresponding control condition. Quantification of tetraspanins release was determined using the same process, except that each condition was corrected by its dilution factor (DF): *BF*DF / (BF*DF + CL*DF)*.

### Detection of molecular complexes by immunoprecipitation

C91-PL and C8166 cells were harvested as explained in the *Western Blot analysis* section. Pellets were lysed using the following buffer: 50mM Tris-HCl pH=7.4, 150mM NaCl, 1mM EDTA, 1mM CaCl2, 1mM MgCl2, 1% CHAPSO, Protease inhibitors (Lysis buffer). 500µg of total proteins were incubated overnight with or without 1µg of anti-tetraspanins antibodies (αCD9, αCD81, αCD82; see supplemental Table S1) on a rotator at 4°C. The remaining lysates were kept and stored at −20°C (*Input*). Then, antibody-protein complexes were deposited on 25µL of protein A magnetic beads (Dynabeads protein A, Life Technologies) and incubated for 4h at 4°C on the rotator. Beads were washed 5 times using 200µL of lysis buffer. Then, the buffer was removed and 20µL of 4X Laemmli buffer was added to the beads. For western blot analysis, samples were heated at 95°C for 10 minutes and loaded onto a 10% SDS-PAGE gel.

### Viral transmission to T-cells

Jurkat reporter T-cells (JKT-LTR-Luc, 50000 cells) were plated in 96-well plates and co-cultured with HTLV-1-infected T-cells (10000 cells) transduced or not (WT) with tetraspanins-specific shRNAs or non-transduced control cells (Jurkat, 10000 cells) for 24h. Cell viability was measured using trypan blue exclusion counting. Reporter cells’ luciferase activity was determined using the luciferase reporter assay system (Promega) following the manufacturer’s instructions. Luciferase activity was measured using a 96 microplate luminometer (Mithras or GloMax), and the background signal due to leaky luciferase activity of Jurkat-LTR-Luc cells was subtracted. Results were presented as normalized to the control (JKT-LTR-Luc co-cultured with WT C91-PL cells) in %.

### NGI-1 drug treatment

C91-PL cells (500000 cells) were seeded in 12-well plates and incubated with or without different concentrations of N-linked Glycosylation Inhibitors 1 (NGI-1) for 48 hours at 37°C. Concentrations of NGI-1 tested: 0µM (DMSO only), 0.5µM, 5µM, or 50µM. Following drug treatment, cell viability was measured using trypan blue exclusion counting. Cells were then collected, washed, and resuspended in 1mL of 1X PBS: 500µL of cells were fixed and plated on 0.01% poly-L-lysine (Sigma-Aldrich) coated glass-bottom Fluorodishes (WPI) for immunofluorescence analysis, and the remaining 500µL were resuspended in 60µL of RIPA lysis buffer for western blotting (see above).

## Statistical analyses

Statistical tests were performed with Prism software (GraphPad), using Kruskal-Wallis or Ordinary one-way ANOVA tests for multiple comparisons. Significant differences were represented as follows: ns = p.value > 0,05; * = p.value ≤ 0.05; ** = p.value ≤ 0.01; *** = p.value ≤ 0.001 and **** = p.value ≤ 0.0001.

## ACKNOWLEDGMENTS

The authors would like to thank Marie-Pierre Blanchard (MRI, CNRS Montpellier, France) for STED microscopy training and Frederic Eghaian (Abberior, Germany) for providing all STED-compatible secondary antibodies. We are also very grateful for Sebastien Lyonnais’ (CEMIPAI, CNRS Montpellier, France) expertise and feedback on AFM imaging. Mathilde Decourcelle and Martial Seveno from the Functional Proteomics Platform of Montpellier (France) are thanked for the mass spectrometry analysis. We acknowledge the Montpellier Imaging Center for Microscopy (MRI) and the CEMIPAI BSL-3 facility for providing excellent working conditions. We thank Cyril Favard (IRIM, CNRS Montpellier, France) for his input on automated image analysis (in Figure 5). The illustrations were created with BioRender.com. This work was supported by the French Agency for Research on AIDS and Viral Hepatitis (grant ANRS0016); institutional funds from the Centre National de la Recherche Scientifique (CNRS); and a 3-year CBS2 Ph.D. fellowship from Montpellier University (UM, France).

## AUTHOR CONTRIBUTIONS

CA performed cell culture, plasmids cloning, cells electroporation, isolation of viral biofilms, lentivectors production, shRNAs transduction, samples preparation for biochemistry and microscopy, western blot assays, confocal and STED images acquisition, AFM imaging, living cells assay, image analysis. HD provided all HTLV-1 infected cell lines, HTLV-1 targeting antibodies, and shCD82-carrying plasmids. SM performed shRNAs transductions for transmission assays, co-culture experiments, and luciferase measurements. PR and JBG performed electron microscopy and immuno-electron microscopy sample preparation and imaging. CA wrote the original draft and conceived the figures. DM, HD, PR, and MIT edited the figures and the manuscript. DM, PR, and HD raised funding for the study. DM and HD directed the study.

## Supplemental Video S1

**<u>Video S1: Polarization of Gag-YFP+ biofilms in living chronically infected T-cells, related to</u> Figure 1**. Time-lapse microscopy imaging of living C91-PL cells electroporated with HTLV-1 Gag-YFP, 20 hours post-electroporation. Sequential epifluorescence images were generated every 15min for 5 hours. Arrows indicate pre-formed Gag-YFP(+) clusters that polarize toward the cell-to-cell junction. Scale bar = 10µm.

