## Supplemental Figures & Legends for "Tetraspanin CD82 maintains HTLV-1 biofilm polarization and is required for efficient viral transmission"

| Application | Target (Conjugate) | Species | Manufacturer (Clone, Ref) | Initial concentration | Dilution |
|---|---|---|---|---|---|
| CONFOCAL MICROSCOPY | CD9 | Mouse | BMA Biomedicals (K41, T-1207) | 200 µg/ml | 1/200 |
| | CD81 | Mouse | Santa-Cruz biotech. (1.3.3.22, sc-7637) | 200 µg/ml | 1/200 |
| | CD82 | Mouse | OriGene (C33, AM26701PU-N) | 1000 µg/ml | 1/1000 |
| | CD43 | Rabbit | Thermofisher (SP55, MA5-16339) | 100 µg/ml | 1/100 |
| | HTLV-1 Gagp19 | Mouse | Zeptometrix (45/6.11.1.3, 801108) | 1000 µg/ml | 1/500 |
| | HTLV-1 Envgp46 | Mouse | Zeptometrix (67/5.5.13.1, 0801127) | 1000 µg/ml | 1/1000 |
| | anti-HTLV-1 serum | Human | Helene Dutartre's lab (Patient 1215) | / | 1/500 |
| | Mouse IgG (AF647) | Goat | Thermofisher (polyclonal, A-21235) | 2 mg/mL | 1/2000 |
| | Rabbit IgG (Atto647N) | Goat | Sigma-Aldrich (polyclonal, 40839) | 1 mg/mL | 1/1000 |
| | WGA lectin (AF647) | / | Thermofisher (W32466) | 1 mg/mL | 1/200 |
| STED MICROSCOPY | CD9 | Mouse | BMA Biomedicals (K41, T-1207) | 200 µg/ml | 1/50 |
| | CD81 | Mouse | Santa-Cruz biotech. (1.3.3.22, sc-7637) | 200 µg/ml | 1/50 |
| | CD82 | Mouse | OriGene (C33, AM26701PU-N) | 1000 µg/ml | 1/250 |
| | CD43 | Rabbit | Thermofisher (SP55, MA5-16339) | 100 µg/ml | 1/50 |
| | YFP | Rabbit | Thermofisher (polyclonal, A-11122) | 2000 µg/ml | 1/200 |
| | HTLV-1 Envgp46 | Mouse | Zeptometrix (67/5.5.13.1, 0801127) | 1000 µg/ml | 1/100 |
| | F-actin (STAR Red) | / | Abberior (STAR 635, phalloidin) | 20 µg/ml | 1/100 |
| | Mouse IgG (STAR | Goat | Abberior (polyclonal, 52283) | 1000 µg/ml | 1/100 |
| | Mouse IgG (STAR 580) | Goat | Abberior (polyclonal, 52403) | 1000 µg/ml | 1/100 |
| | Rabbit IgG (STAR Red) | Goat | Abberior (polyclonal, 41699) | 1000 µg/ml | 1/100 |
| | Rabbit IgG (STAR 580) | Goat | Abberior (polyclonal, 41367) | 1000 µg/ml | 1/100 |
| WESTERN BLOT | CD9 | Mouse | BMA Biomedicals (K41, T-1207) | 200 µg/ml | 1/500 |
| | CD81 | Mouse | Santa-Cruz biotech. (5A6, sc-23962) | 200 µg/ml | 1/500 |
| | CD82 | Mouse | OriGene (C33, AM26701PU-N) | 1000 µg/ml | 1/1500 |
| | HTLV-1 Gagp19 | Mouse | Zeptometrix (45/6.11.1.3, 0801108) | 1000 µg/ml | 1/1000 |
| | GAPDH (HRP) | Mouse | Sigma (GAPDH-71.1, G9295) | 1000 µg/ml | 1/25.000 |
| | Mouse IgG (HRP) | Rabbit | Dako (polyclonal, P0260) | / | 1/3000 |

| Application | Target | Sequence |
|---|---|---|
| shRNAs for TRANSDUCTIONS | None (shCtrl) | 5'– GATCCCCGACCCCCTTGTGAATCTCTTCAAGAGAGAGAT-TCACAAGGGGGTCTTTTTGGAAA-3 |
| | CD9 | 5'-CCGGCACAAGGATGAGGTGATTAAGCTCGAGCTTAATCACCTCATCCTTGTGTTTTTG-3' |
| | CD81 (L35) | 5'-GATCCCCCGCTGTCATGATGTTCGTTTCAAGAGAACGAACATCATGACAGCGTTTTT-GGAAA-3' |
| | CD81 (L36) | 5'-GATCCCCTCATGATGTTCGTTGGCTTTCAAGAGAAGCCAACGAACATCATGATTTTT-GGAAA-3' |
| | CD82 (32) | 5'-CCGGCTTCTACAACTGGACAGACAACTCGAGTTGTCTGTCCAGTTGTAGAAGTTTTTG-3' |
| | CD82 (33) | 5'– CCGGGTTTCATCTCTGTCCTGCAAACTCGAGTTTGCAGGACAGA-GATGAAACTTTTTG-3' |

**Table S1: Antibodies and shRNAs used in this study.** Table gathering all the antibodies used in this study: Primary antibodies (blue) and secondary antibodies (orange) working dilutions are provided. Also, sequences of shRNAs used for transduction experiments are given (red).

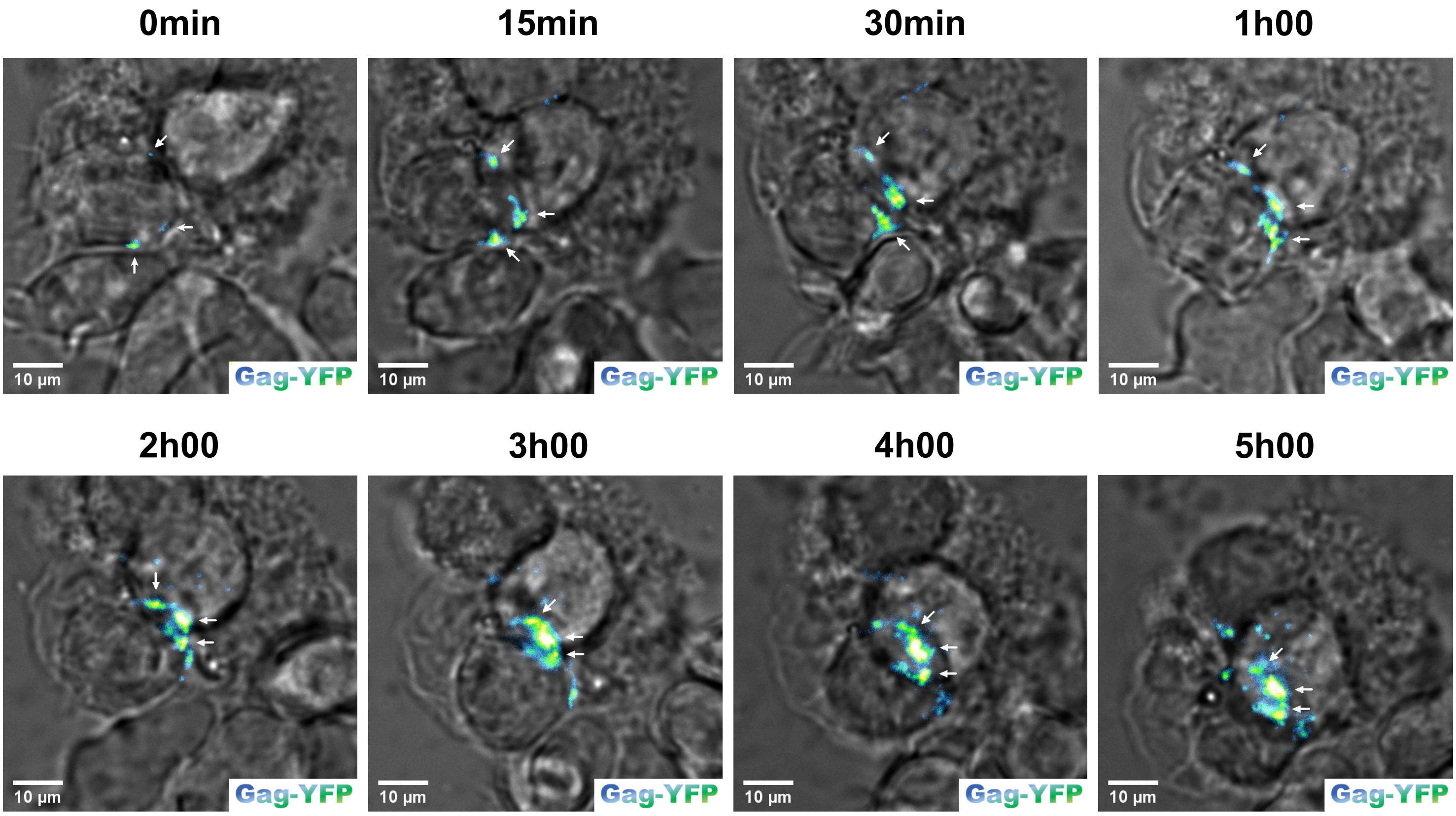


**Figure S1: Time-lapse of Gag-YFP+ biofilms formation in living chronically infected T-cells, related to Video S1 and Figure 1.** Time-lapse of Gag-YFP+ biofilms formation: Living C91-PL cells were electroporated with HTLV-1 Gag-YFP and imaged 20h post-electroporation using an automated confocal laser-scanning microscope at 37°C, 5% CO2. Epifluorescence images were generated every 15min for 5 hours. White arrows point toward pre-formed Gag-YFP+ clusters (in green fire blue) that translocate toward the intercellular contact. Scale bars = 10μm.

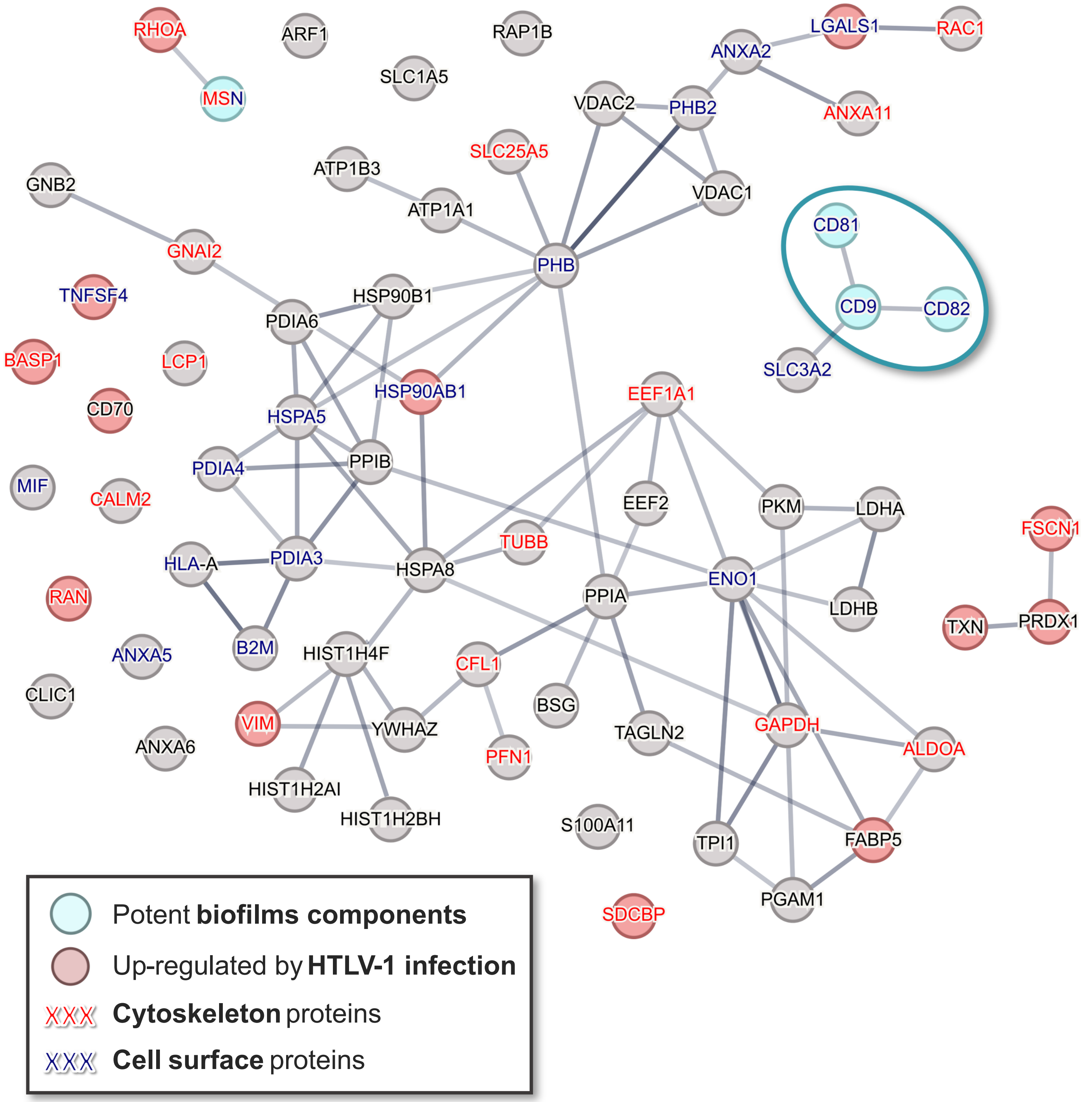


**Figure S2: STRING interaction network of cellular components identified in isolated HTLV-1 biofilms by mass spectrometry, related to Figure 2B.** STRING protein–protein association network of 69 cellular components identified with high confidence in C91-PL biofilms-enriched supernatant (n = 3). To select these proteins, contaminants were removed, only the 100 most abundant proteins were kept (based on the IBAQ score), and molecules identified with less than 3 peptides were excluded. Gray lines indicate that the proteins are part of a physical complex (interaction score of medium confidence from experimental data). Blue circles highlight potent biofilm components; Red circles indicate proteins up-regulated by HTLV-1 infection (based on mass spectrometry analyses of infected primary T-cells compared to non-infected T-cells, GSE17718 database, Kress et al, 2010). Red or blue text indicates cytoskeleton or cell surface proteins, respectively (Cellular Component Gene Ontology).

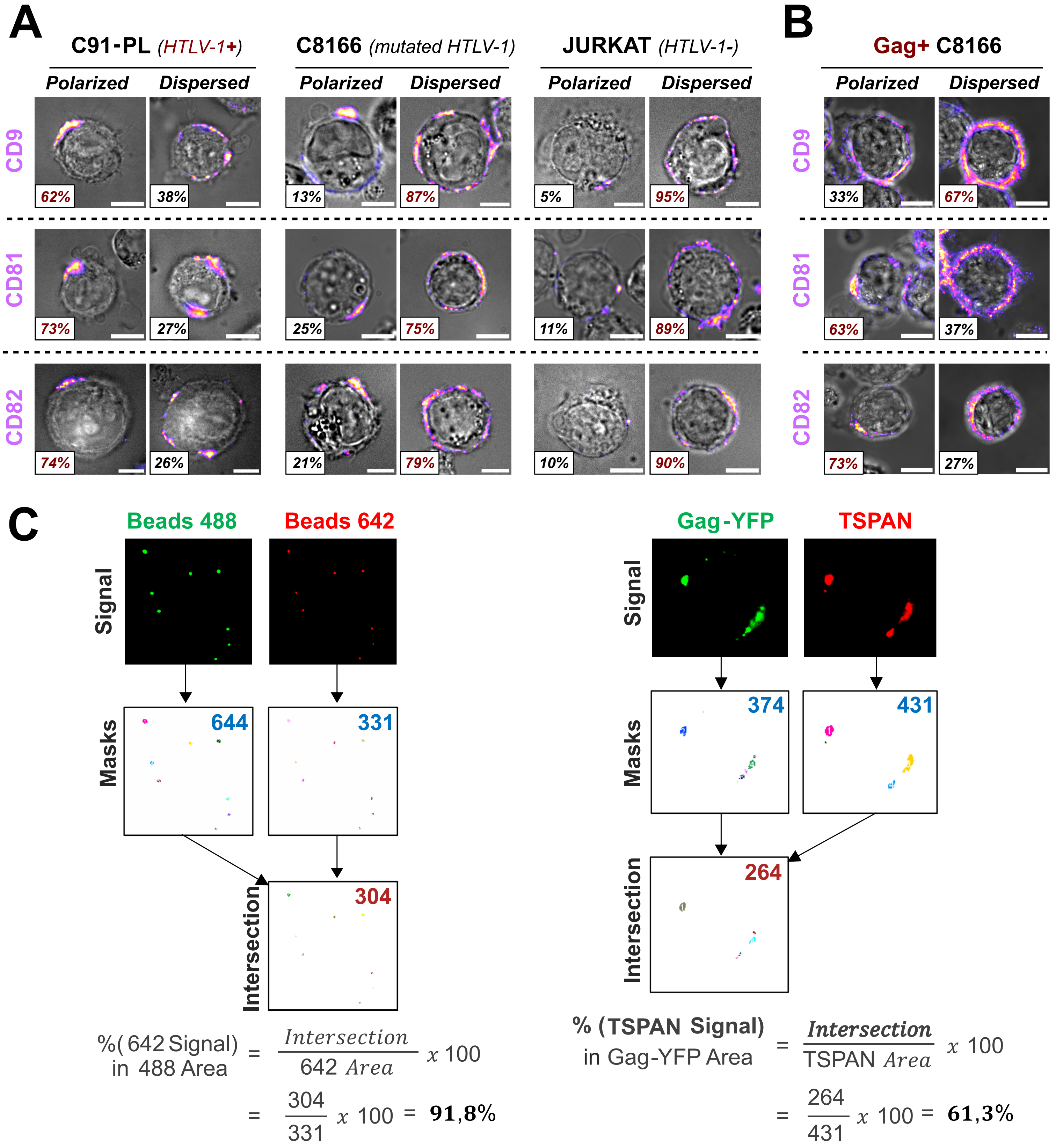


**Figure S3: HTLV-1 Gag is sufficient to initiate the polarization of tetraspanins at the surface of T-cells, related to Figure 3. (A)** Representative images of CD9, CD81, and CD82 expression patterns (fire red) at the surface of C91-PL, C8166, or Jurkat T-cells: *"Polarized"* or *"Dispersed"*. **(B)** Representative images of CD9, CD81, and CD82 organization at the surface of C8166 cells electroporated with HTLV-1 Gag. Percentages indicate the proportion of cells displaying each phenotype (red = predominant one). Scale bars = 10µm. (**C**) Colocalization analysis between tetraspanins and Gag-YFP+ biofilms, related to Figure 3F. Left: Measurement of the overlapping between green and red channels using dual-color beads to assess instrumental error. Right: % of tetraspanin (TSPAN) total area (red) overlapping with Gag-YFP area (green). In both panels, thresholding was performed to select only the higher pixel intensities. Masks obtained following thresholding were used to calculate the total area of the signals for each channel (blue) and their intersection (red) in µm2. % of total TSPAN area contained in Gag-YFP area was calculated using the formula examplified bellow.

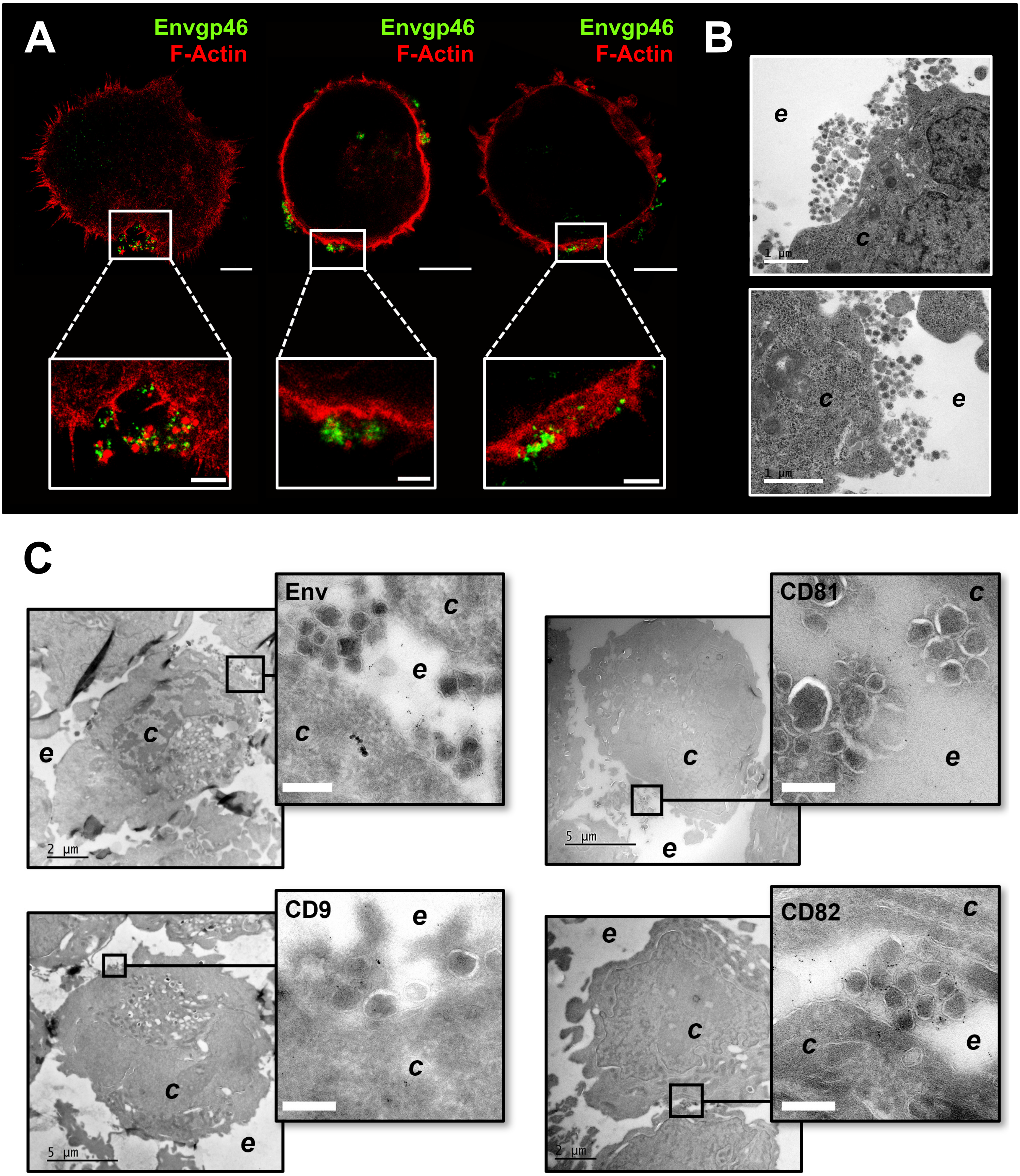


**Figure S4: Organization of HTLV-1 biofilms at the surface of infected T-cells and incorporation of CD9, CD81, and CD82 into viral particles, related to Figure 4A and 4E. (A)** Top: STED 2D images of HTLV-1 chronically infected T-cells (C91-PL) stained for cortical actin (F-actin in red) which highlights cell boundaries, and for HTLV-1 viral envelope (green). Scale bars = 5µm. Bottom: Magnified images showing viral aggregates in membrane folds (left), or on top of cell membranes (middle and right). Scale bars = 1µm. **(B)** Transmission electron microscopy images showing HTLV-1 biofilms at the surface of C91-PL cells. Scale bars = 1µm. **(C)** Immuno-electron microscopy images of HTLV-1 biofilms at the surface of C91-PL cells stained for Envgp46, CD9, CD81, or CD82 with antibodies coupled to 6nm gold beads (black spots). *c = cytosol ; e = extracellular environment.* White scale bars = 200nm.

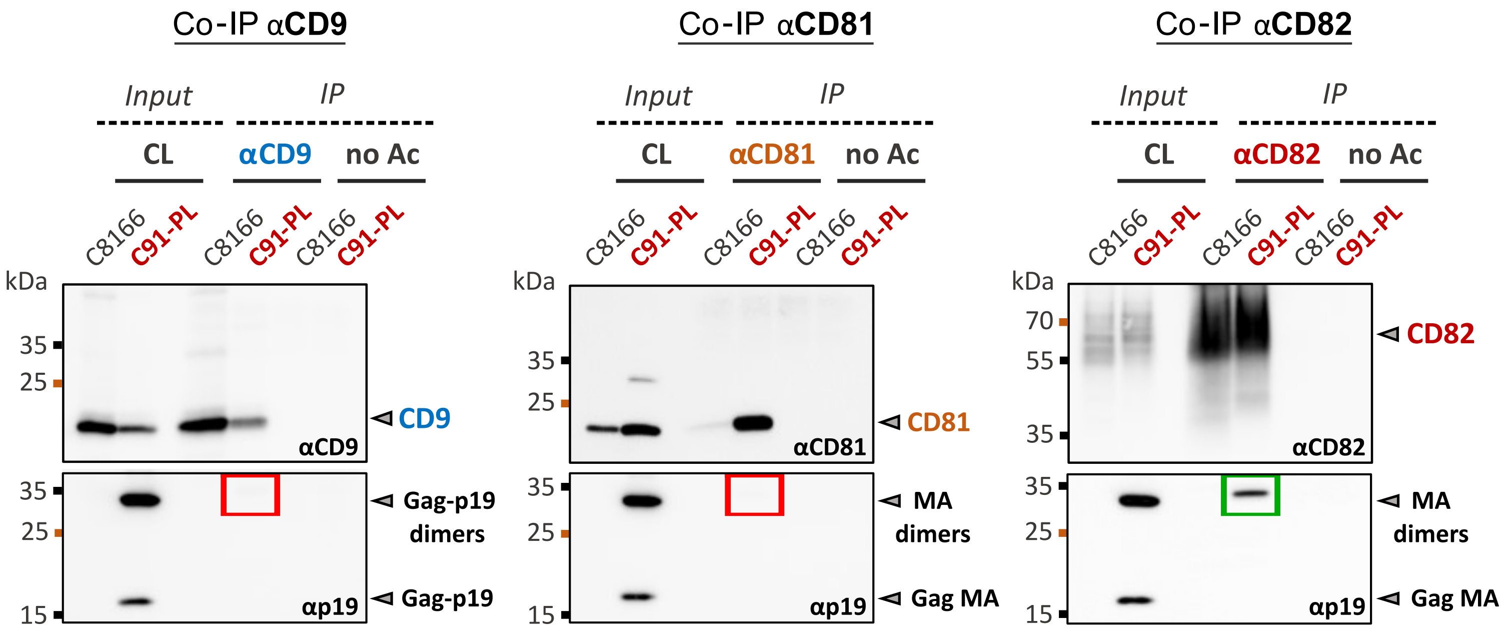


**Figure S5: CD82 forms a complex with HTLV-1 Gag, related to Figure 5.** Representative immunoblots of three independent co-immunoprecipitation experiments: *Input* = C91-PL or C8166 cell extracts were directly immunoblotted with anti-HTLV-1 Gag-p19 and anti-tetraspanin antibodies. *IP* = Cell extracts were immunoprecipitated with αCD9, αCD81, or αCD82 antibodies and blotted with anti-HTLV-1 Gag-p19 or anti-tetraspanin antibodies. CD9 = 24kDa, CD81 = 25kDa, CD82 = 45-60 kDa, Gagp19 = 19kDa. Bands at 35kDa are dimers of Gag-p19 preserved in non-reducing conditions.

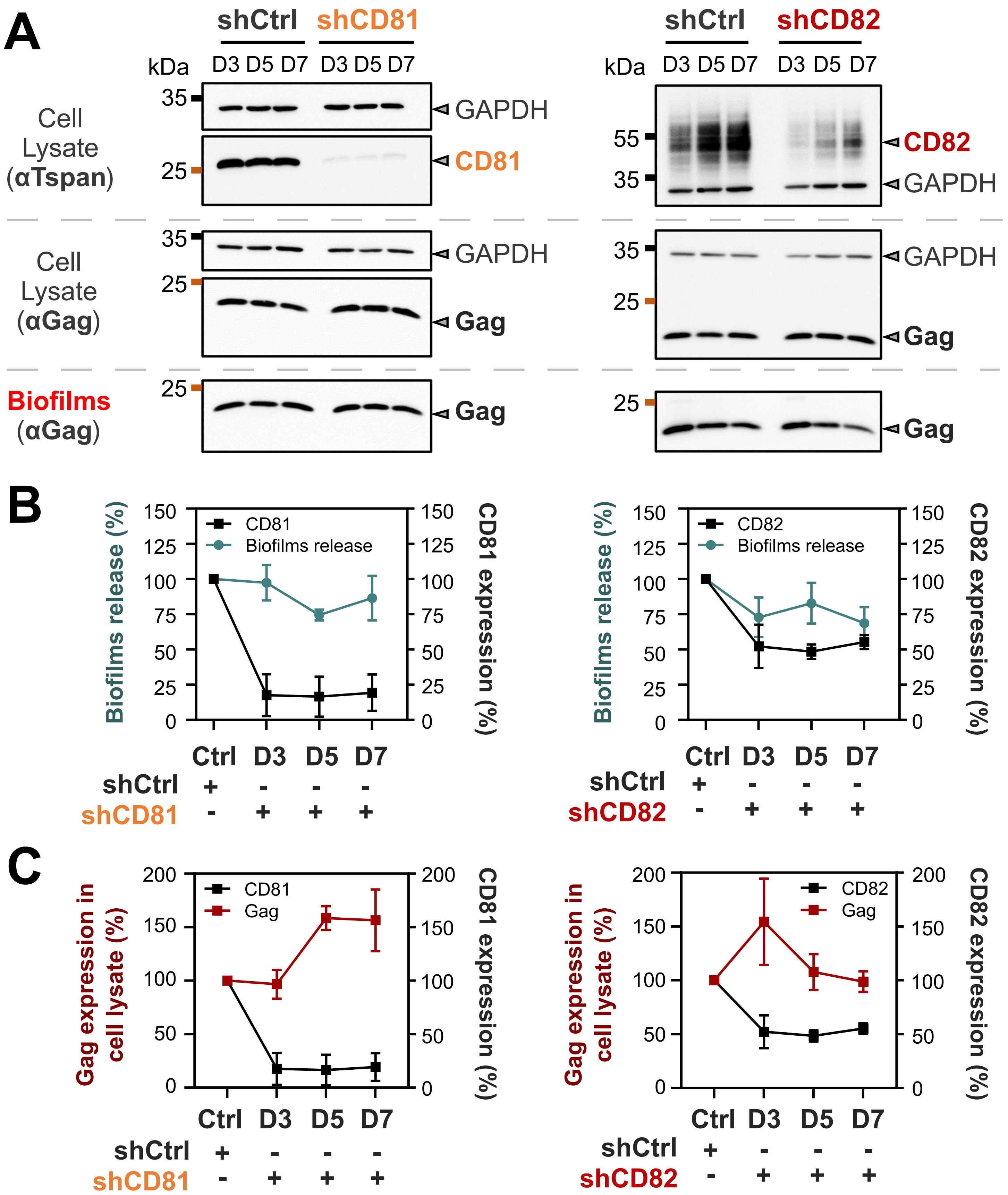


**Figure S6: Impact of CD81 and CD82 shRNA silencing on HTLV-1 biofilms release, related to Figure 6C. (A)** Effects of CD81 (left) and CD82 (right) shRNA-silencing on the release of HTLV-1 biofilms from C91-PL cells, assessed by Western-Blot at three different time-points post-transduction (Day 3, Day 5, and Day 7). On the immunoblots, Gagp19 = 19kDa, CD81 = 26kDa, CD82 = 45-60 kDa. CD82 is highly glycosylated as shown by the multiple bands around 55kDa. Endogenous GAPDH is used as a loading control for cell lysates and Gagp19 is used to stain viruses. **(B)** Graphs showing biofilms' release in % (green curves) calculated as follows: (Gagp19 BF / (Gagp19 BF + Gagp19 CL))*100 as compared to the control (shCtrl) which was normalized to 100% for each time point. CD81 or CD82 silencing is represented by the black curves. CL = Cell lysate. BF = Biofilms. **(C)** Graphs showing the % of Gag expression in the cell lysate for all conditions. n = 3 independent experiments.

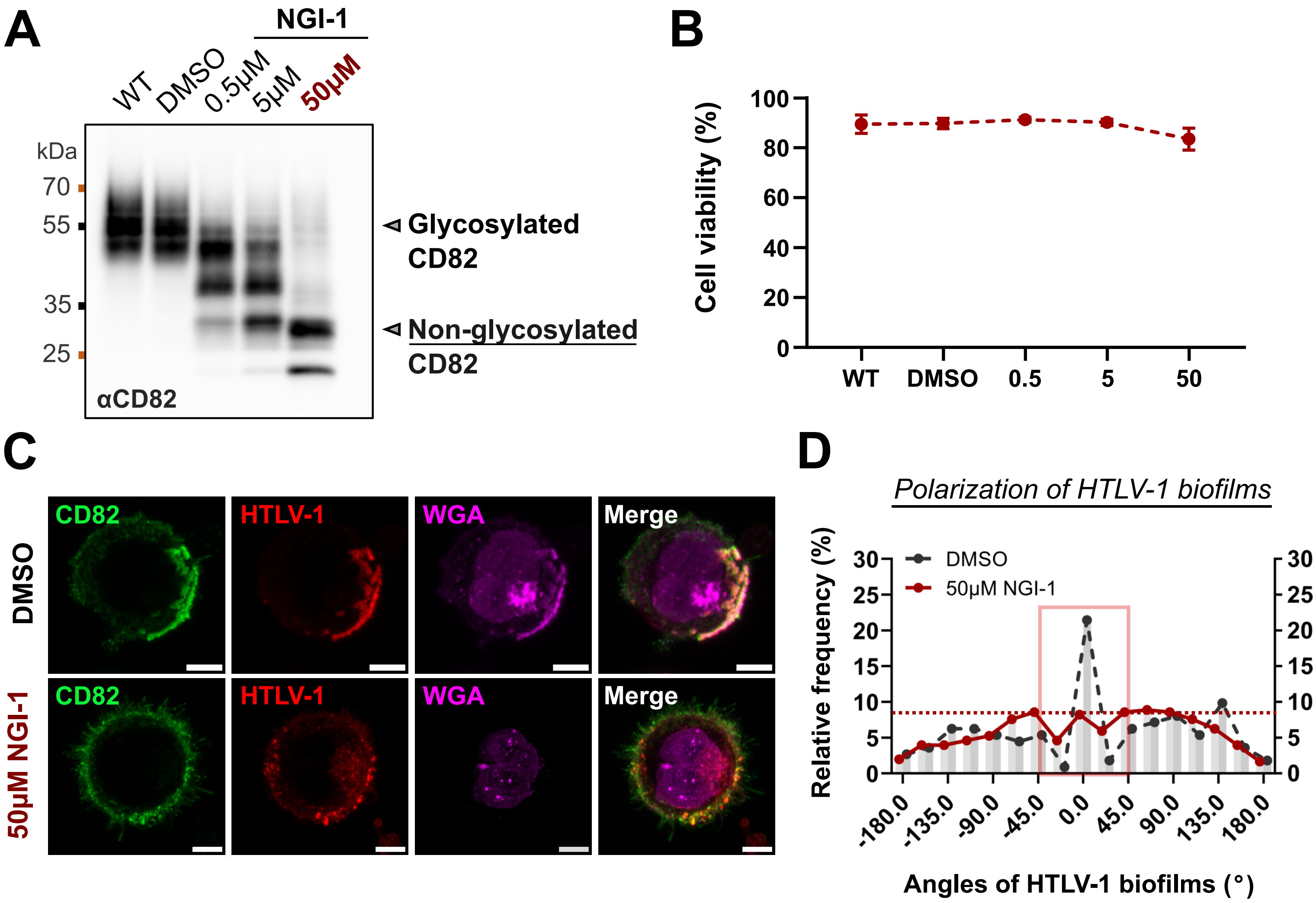


**Figure S7: N-glycosylation is a key parameter for HTLV-1 biofilm polarization. (A)** Representative immunoblot showing CD82 glycosylation profiles in C91-PL cells treated or not (WT, DMSO) with different concentrations of NGI-1 (0.5µM, 5µM, or 50µM) for 48 hours. **(B)** Viability of C91-PL cells ± SD (%) related to panel A. n = three independent experiments. **(C)** Representative confocal images of C91-PL cells treated or not with NGI-1 as in panel A. All cells were fixed and stained for CD82 (green), for HTLV-1 using serum from a HAMP-TSP patient (magenta), or for glycoproteins using WGA-conjugated lectins (magenta). Scale bars = 5µm. **(D)** Frequency distribution of HTLV-1 biofilms angles at the surface of C91-PL cells treated or not with NGI-1 as in panel A (n = 25 cells; n > 140 HTLV-1 clusters per condition). While a peak between -45° and 45° (red square) is representative of a polarized profile, the absence of a peak indicates a depolarized pattern (red dotted line).
